# Cleavage site heterogeneity at the pre-mRNA 3′-untranslated region regulates gene expression in oxidative stress response

**DOI:** 10.1101/2024.09.11.612397

**Authors:** Feba Shaji, Jamshaid Ali, Rakesh S. Laishram

**Author notes:** Corresponding Address: Rakesh S. Laishram, Ph.D., Rajiv Gandhi Centre for Biotechnology, Melaranoor Road, Jagathy, Trivandrum, Kerala, 695014, India, E mail.

## Abstract

Endonucleolytic cleavage step of the eukaryotic mRNA 3′-end processing is considered imprecise that leads to heterogeneity of cleavage site (CS) with hitherto unknown function. Contrary to the popular belief, we show that this imprecision in cleavage is tightly regulated resulting in CS heterogeneity (CSH) that controls gene expression in antioxidant response. CSH centres at a primary CS followed by a number of subsidiary cleavages that is determined by the position of the CS. Globally and by using reporter antioxidant NQO1 mRNA, we discovered an inverse relationship between the number of CS and the gene expression with the primary CS exhibiting highest cleavage efficiency. Strikingly, reducing CSH and increasing primary CS usage induces gene expression. Under oxidative stress condition (tBHQ, H_2_O_2_ or NaAsO_2_), there is a decrease in the CSH and an increase in the primary CS usage to induce antioxidant gene expression. Concomitantly, ectopic NQO1 expression from the primary CS or reduction in CSH imparts cellular oxidative stress tolerance. Genome-wide CS analysis of stress response genes also shows a concomitant result. We show that oxidative stress induces affinity/strength of cleavage complex assembly increasing the fidelity of cleavage at the primary CS thereby reducing CSH inducing antioxidant response.

## INTRODUCTION

Processing at the 3′-end of a precursor messenger RNA (pre-mRNA) plays a pivotal role in gene expression(Boreikaite & Passmore, 2023; Neve *et al*, 2017). The processing reaction involves two steps (cleavage and polyadenylation, CPA) that requires cleavage and polyadenylation specificity factor (subunits CPSF160, WDR33, CPSF100, hFIP1, CPSF73, and CPSF30), cleavage stimulating factor (subunits CstF77, CstF64, and CstF50), cleavage factor (CFIm and CFIIm), scaffolding protein symplekin, poly(A) binding protein (PABPN1) and poly(A) polymerase (PAP) as core components(Boreikaite & Passmore, 2023; Mandel *et al*, 2008; Xiang *et al*, 2014). While CPSF, CstF, PAP, CFIm and IIm are involved in the cleavage reaction, polyadenylation requires PAP, CPSF and PABP(Bienroth *et al*, 1993; Gilmartin & Nevins, 1991; Kuhn *et al*, 2009; Takagaki *et al*, 1989). CPSF subunits WDR33 and CPSF30 recognises the poly(A) signal (PA-signal) hexamer (AAUAAA) to assemble the CPA complex at the poly(A) site (PA-site)(Chan *et al*, 2014; Schönemann *et al*, 2014; Sun *et al*, 2018). CPSF160 cooperates with CstF64 that binds the downstream sequence element (DSE, GU/U-rich ∼15-30 nucleotides downstream of the PA-signal)(Gilmartin & Nevins, 1991; MacDonald *et al*, 1994; Murthy & Manley, 1995). Along with CFIm and CFIIm and other core components, CPSF assembles the CPA complex(Brown & Gilmartin, 2003; de Vries *et al*, 2000; Ruegsegger *et al*, 1996). CPSF73 then cleaves the pre-mRNA at the cleavage site (CS) followed by PAP-mediated PA-tail addition and PABPN1 coating of the PA-tail(Deo *et al*, 1999; Mandel *et al*, 2006; Raabe *et al*, 1994; Ryan *et al*, 2004). Canonical PAPα/γ is the primary PAP for mRNA polyadenylation in the nucleus(Kyriakopoulou *et al*, 2001; Raabe *et al*, 1991; Topalian *et al*, 2001; Wahle *et al*, 1991). The discovery of a variant PAP, Star-PAP has revealed existence of alternative PAPs for nuclear polyadenylation(Laishram & Anderson, 2010; Mellman *et al*, 2008).

CPA has a primary role in gene expression primarily through 1. control of mRNA metabolism and translation, 2. co-ordination with transcription and other RNA processing events, and 3. alternative polyadenylation(Boreikaite & Passmore, 2023; Elkon *et al*, 2013; Komili & Silver, 2008; Kumar *et al*, 2019; Moore & Proudfoot, 2009; Passmore & Coller, 2022). The PA-tail serves at least three functions: first, PA-tail through its coating with PABP protects the mRNA from 3′- to 5′-exonucleases along with the 5′-end m7G cap; second, it serves as the initiation site for mRNA turnover by deadenylation; and third, PA-tail shortening precedes decapping for mRNA decay(Mitchell & Tollervey, 2000; Passmore & Coller, 2022; Weill *et al*, 2012). In addition to the stability, PA-tail coating by PABP facilitates mRNA nuclear exit and also acts as enhancers of mRNA translation initiation by association with transcription initiation factors(Imataka *et al*, 1998; Kahvejian *et al*, 2005; Qu *et al*, 2009; Shi *et al*, 2017). Additionally, many of the core CPA components are detected at the promoter along with PolII and also couple with cap binding protein forming a closed loop to coordinate with transcription initiation(Dantonel *et al*, 1997; Flaherty *et al*, 1997; Jiao *et al*, 2013; Mapendano *et al*, 2010). Furthermore, a number of CPA components assembles with spliceosomal components which stimulates intron splicing(Awasthi & Alwine, 2003; Huang *et al*, 2023; Kyburz *et al*, 2006; Millevoi *et al*, 2006; Vagner *et al*, 2000). Apart from these mechanisms, CPA also regulates gene expression through (alternative polyadenylation) APA where a gene encodes for more than one transcript with different length due to the presence of more than one PA-site(Di Giammartino *et al*, 2011; Mohanan *et al*, 2022; Tian & Manley, 2017). APA alters the primary structure of a protein, or transcript length affecting stability, translation, localisation and encoded protein interaction(Di Giammartino *et al*., 2011; Mohanan *et al*., 2022; Tian & Manley, 2017).

While cleavage and polyadenylation are two coupled steps in the 3′-end processing, cleavage step is considered imprecise that typically occurs within 10-25 nucleotides downstream the PA-signal (Chen *et al*, 1995; Weiss *et al*, 1991). The positioning of the PA-signal and the DSE limits the region where cleavage can occur (Weiss *et al*., 1991). Nevertheless, the predominant nucleotide at the CS appears as adenosine followed by uridine, cytosine, and guanosine (A>U>C>>G) and the dinucleotide composition at the CS (CA>AA>TA>>GA) in the order(Chen *et al*., 1995). Intriguingly, the imprecision in the cleavage leads to multiple cleavage events instead of cleaving at only one CS between the PA-signal and the DSE resulting in cleavage site heterogeneity (CSH) (Chen *et al*., 1995; Pauws *et al*, 2001; Stroup & Ji, 2023). Unlike in APA where more than one PA-sites are alternatively used at the 3′-UTR that leads to distinct mRNA isoforms with different UTR lengths, in CSH, cleavage occurs at multiple nucleotide sequences between the PA-signal hexamer and the DSE of a single PA-site (Chen *et al*., 1995; Stroup & Ji, 2023)(Schematics to clarify CSH, APA and PA-site is shown in Supplementary Fig. 1A). CSH neither leads to discernible difference in the UTR length nor generate significant mRNA isoforms(Chen *et al*., 1995; Pauws *et al*., 2001; Stroup & Ji, 2023). It was considered stochastic and its mechanism, regulation or cellular ramifications of the imprecision in cleavage is unclear. Nevertheless, both gain and loss of function mutations of CS are reported in various diseases(Bishop *et al*, 1988; Gehring *et al*, 2001; Sheets *et al*, 1990). Therefore, we investigated the cellular and mechanistic implications of CSH in gene expression.

Global analysis shows CSH happens with a tight regulation wherein cleavage centers at a primary CS followed by a number of subsidiary CS. This CSH is linked with the fidelity of cleavage such that mutation of the subsidiary CS increases cleavage at the primary CS. Global and mutational analysis of a reporter antioxidant NQO1 mRNA, we discovered an inverse relationship between the number of CS and the gene expression. The primary CS shows highest cleavage efficiency such that reducing CSH and increasing primary CS usage induces gene expression. Cleavage at the primary CS and hence cleavage efficiency is determined by the position of the CS downstream of the PA-signal and not by the sequence composition. This mechanism regulates oxidative stress response where increased antioxidant protein expression involves both reduction in the CSH and an enhanced primary CS usage. Consistently, in the cellular assays, cells expressing ectopic NQO1 with reduced CSH or with the primary CS imparts cellular tolerance to H_2_O_2_ and NaAsO_2_ stress. We show that oxidative stress induces affinity/strength of cleavage complex assembly increasing the fidelity of cleavage at the primary CS thereby reducing CSH inducing antioxidant response. To our knowledge, this is the first example of a biological significance of CS imprecision or heterogeneity that regulates gene expression in antioxidant response.

## RESULTS

### Global cleavage site analysis reveals existence of cleavage site heterogeneity in majority of the cellular mRNAs covering nearly half of the PA-sites genome wide

To understand the regulation and ramifications of imprecisions in the endonucleolytic cleavage step leading to CSH in gene expression, we first analysed the global 3′-READS (3′-Region Extraction and Deep Sequencing) data (Accession Number: GSE84461) for the CS selection in each PA-site genome-wide in HEK293 cells (Li *et al*, 2017). We define a PA-site as a region at the 3′-UTR where cleavage and polyadenylation occurs encompassing the PA-signal and other cis-elements required for CPA reaction; CS is defined as an actual nucleotide where cleavage occurs and PA-tail is added such that a transcript from a particular a PA-site can have either one or more CS (**Supplemental Fig. 1A**). For clarity, schematics of CSH and comparison with that of APA is shown in **Fig. 1A and Supplemental Fig. 1A**. In CSH, cleavage of a PA-site occurs at more than one CS within 10-30 nucleotides downstream of the AAUAAA PA-signal resulting in heterogenous cleavages such that a single PA-site exhibits multiple CS with no significant difference in the apparent mRNA length. Whereas, in APA, different PA-sites at the 3′-UTR are alternatively used that leads to distinct mRNA isoforms with difference in the UTR lengths (Mohanan *et al*., 2022). Each PA-site in APA can exhibit CSH. In our analysis, we obtained a total of ∼800000 processed reads aligning to distinct PA-site(s) on respective CS of assigned genes. Altogether, there were >175,000 CS clustered across >70,000 PA-sites on 12,000 genes indicating the existence of CSH globally.

**Figure 1:**
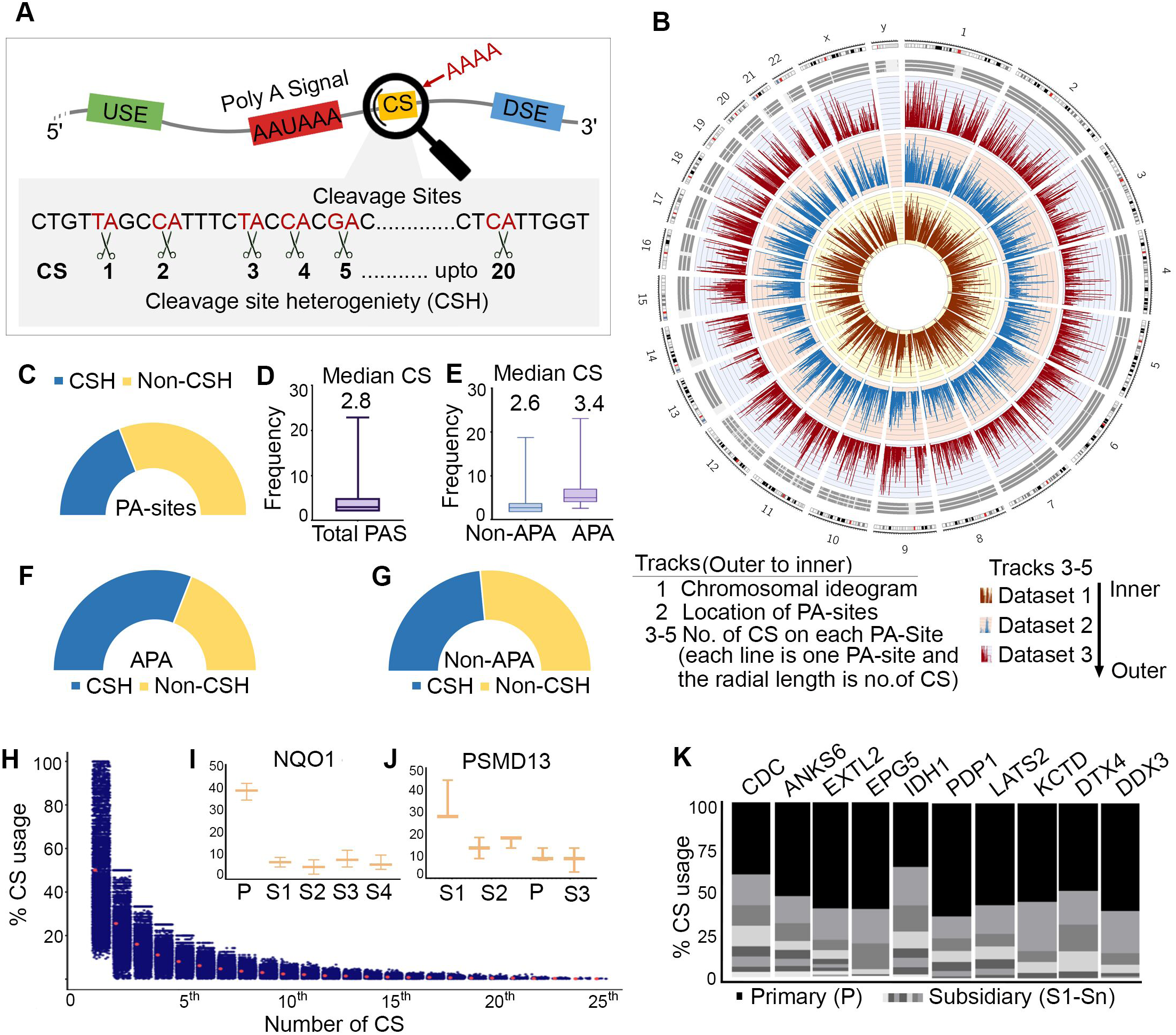
Global cleavage site analysis reveals heterogeneity in majority of the cellular mRNAs covering nearly half of the PA-sites genome wide. (*A*) Schematics describing cleavage site heterogeneity (CSH) of a particular PA-site. Various cis elements in the 3′-UTR of RNA (PA-signal, downstream sequence element, and cleavage sites are indicated). The zoomed-in view of various CS showing CSH is represented. (*B*) Circos plot to visualise the genome-wide distribution of CSH of each PA-sites plotted against the chromosomal location and to assess the overlap of CS across of 3′-READS datasets. The outermost track of the plot represents the chromosome ideogram showing the physical location of each gene. The three inner tracks correspond to the cleavage sites of different mRNAs that has been plotted against exact genomic location from three different datasets. The radial length of each track corresponds to the number of CS thus representing heterogeneity. The circumference (X axis) represents the location of the PA-site and the radial length (Y axis) represents the number of CS of each PA-site. (*C*) Half donut plot for % of total PA-sites that show CSH (blue) versus that do not show (yellow). (*D*) Box plot of frequency of CSH of PA-sites genome wide. Median CS of PA-sites is indicated on the top. (*E*) Box plot of frequency of CSH of PA-sites from APA-dependent and -independent mRNAs. Median CS of PA-sites from APA-dependent and-independent mRNAs is indicated. (*F*) Half donut plot of PA-sites from the mRNAs that are APA-dependent, blue and yellow colours represent % PA-sites with CSH and non-CSH respectively. (*G*) Half donut plot of PA-sites from the mRNAs that are non-APA regulated mRNAs (blue and yellow colours represent % PA-sites with CSH and non-CSH respectively). (*H*) Dot plot showing distribution of % usage of different CS (numbered from 1^st^ to 25^th^ CS). The CS that is used maximally is assigned 1^st^ (primary CS) followed by CS numbers based on the rank of CS usage (2^nd^ to 25^th^) (subsidiary CS). (*I-J*) Box plot of % usage of different CS of *NQO1* (chromosomal locations of each CS are P: 69709403, S1: 69709401, S2: 69709407, S3: 69709409 and S4: 69709417) and *PSMD13* mRNA (chromosomal locations of each CS are P: 252984, S1: 252979, S2: 252980, S3: 252981 and S4: 252982). (*K*) Stacked column plot of % usage of primary versus subsidiary CS of 10 select mRNAs (*CDC, ANKS6, EXTL2, EPG5, IDH1, PDP1, LATS2, KCTD, DTX4, DDX3*). Primary CS is indicated in black and different subsidiary CS are indicated in different shades of grey.

Our analysis revealed two classes of PA-sites on mRNAs, first having only one CS, and the other on majority of mRNAs having more than one CS (that confirms the occurrence of CSH). Circos plot of three experimental replicates showed genome wide presence of CSH on numerous PA-sites (**Fig. 1B**). The outermost track of the plot represents the chromosome ideogram showing the physical location of each gene on the chromosome. The three inner tracks correspond to the CS of different mRNAs from three different datasets that were plotted against respective genomic locations. The radial length of each track corresponds to the number of CS thus representing heterogeneity. Thus, each line in the plot represents distinct PA-sites and the length of the line corresponds to the number of CS. We observed that the position and length of the lines from various replicates overlapped at the similar chromosomal positions confirming CSH (**Fig. 1A**). CS clusters ranged from 1 to 20 on different PA-sites, where majority of the PA-sites had 3 to 6 CS with a median of ∼2.8 CS per PA-site (**Fig. 1B-C**). The number of CS determines the degree of heterogeneity of cleavage and we observed more PA-sites with low heterogeneity (less number of CS) than that of higher heterogeneity (more number of CS). This is also seen from a jitter plot where each dot represents one PA-site with a particular (n) number of CS (**Supplemental Fig. 1B**). In this plot, each blue dot represents a PA-site in the X-axis that are plotted against the number of CS of that particular PA-site in the Y-axis. The length of X axis is an arbitrary value to increase the visibility of the dot and the density of the dots correspond to number of PA-site. The distribution pattern of the PA-site with different numbers of CS showed a gradual decrease in the number of PA-sites with increasing number of CS (**Supplemental Fig. 1B**). Overall, CSH was observed on 10944 mRNAs (majority of mRNAs) at different PA-sites of which around half (∼45%, 30,500 out of 70,000 PA-sites on different mRNAs) exhibited CSH (**Fig. 1C**). The median value of CSH observed on PA-sites corresponds to 2.8 (**Fig. 1D**). Intriguingly, PA-sites of alternatively polyadenylated mRNAs (APA-dependent) showed more heterogeneity than that of the APA-independent mRNAs (**Fig. 1E**). The median number of CS (cleavage per PA-site) of the APA-dependent mRNAs was 3.4 compared to the median CSH of 2.6 of the non-APA independent mRNAs (**Fig. 1E**). While 60% of the PA-sites on the APA-dependent mRNAs exhibited CSH, only around ∼45% of the PA-sites on non-APA mRNAs showed CSH (**Fig. 1F-G**). An example of CSH on *EXOC3* mRNA showing nucleotide details and the number of CS is shown in **Supplemental Fig. S1C**. Further, we looked into the heterogeneity pattern of target PA-sites of two major PAPs in the nucleus (canonical PAPα and the non-canonical Star-PAP) that polyadenylates distinct target PA-sites. We did not see any significant difference in the CSH pattern between the PA-sites regulated by Star-PAP and that of PAPα (**Supplemental Fig. S1D-E**). The number of median PA-site among the targets of the two PAPs was ∼2.8 indicating a similar CSH pattern between the target PA-sites of the two PAPs. Together, these results reveal a global CSH with a higher CSH of PA-sites of APA-dependent than that of non-APA dependent mRNAs.

### Cleavage site heterogeneity is tightly regulated and exhibits preference of one primary site over other subsidiary sites

Strikingly, among the heterogeneous CS, a single CS was predominantly used resulting in cleavage events at the predominant CS surpassing cleavages at other sites genome-wide (**Fig. 1H**). We refer to this preferential CS as the “primary CS” and the other lesser-used CS of a PA-site as the “subsidiary CS” such that a PA-site can have a primary site and a number of subsidiary sites. Dot plot analysis of CS ranked according to their usage shows a predominance of the primary CS site over other subsidiary sites with a marked decrease in the usage from primary CS to various subsidiary sites (**Fig. 1H**). The distribution of primary CS usage ranged from ∼20% to 99% with a median usage of 65%, whereas the subsidiary CS usage ranged from 1% to 50% with an overall median usage of ∼14% for the subsidiary CS. Analysis of the CSH on individual genes also showed similar observation. For example, on *NQO1* mRNA 3′-UTR, the terminal PA-site has five major CS that were consistently observed, one primary and four subsidiary sites (**Fig. 1I**). The primary CS was cleaved >40% on an average while the cleavage on the other subsidiary sites was <5% each. Similarly, *PSMD13* mRNA showed 40% usage of the primary CS among the total 5 CS (**Fig. 1J**). CSH analysis of 10 select mRNAs (*CDC*, *ANKS6*, *EXTL2*, *EPG5*, *IDH1*, *PDP1*, *LATS2*, *KCTD*, *DTX4*, and *DDX3*) also showed similar preferential cleavage at the primary CS over the subsidiary sites (**Fig. 1K**). Further, we validated our genome-wide data of CSH by 3′-RACE assays of 3 genes (*HMOX1*, *NQO1*, and *PRDX1*) followed by Sanger sequencing of the PA-tailed mRNA (**Supplemental Fig. S1F**). We observed a similar CSH pattern with consistent positions of primary and subsidiary CS in both 3′-RACE and 3′-READS data (**Supplemental Fig. S1G**). Together, these results confirm that CSH is tightly regulated for CS selection in the cell.

### Cleavage site heterogeneity on target PA-sites is closely linked with expression of the corresponding mRNA

To understand the significance of CSH, we analyse the relative gene expression from each CS. We assess the total read counts per million of each CS specific transcripts of a PA-site relative to constitutive *GAPDH* level and expressed over total expression level of the particular mRNA. Remarkably, we observed an inverse co-relation between CSH and gene expression. There was a gradual decrease in the expression with an increase in the number of CS (**Fig. 2A**). There was higher relative expression from mRNA isoforms with lower heterogeneity (<10 CS per PA-site) than that of higher heterogeneity, and the expression from the PA-sites with CS beyond 15 were marginal relative to total mRNA expression (**Fig. 2A**). A plot of average relative expression per PA-site also showed highest expression with mRNAs with lower CSH (2-6 CS), a moderate expression with medium CSH (7-15 CS) and lowest expression with highest CSH (>18 CS) (**Fig. 2B**). Consistently, we observed a higher relative expression from mRNA isoforms with no CSH (having one CS) compared to those mRNA isoforms with multiple CS (**Fig. 2C**). Further, randomly selected mRNA isoforms with different CSH level showed a linear decrease in the expression with the increasing number of CS (**Supplemental Fig. S2A**). Additionally, since primary CS is predominantly used, we then analysed the expression from the primary CS of different PA-sites. The expression from the primary CS of PA-sites having low CSH was higher than that of PA-sites having high CSH, and a gradual decrease in the expression was observed with increasing number of CS (**Fig. 2D**). Analysis of individual genes (*NQO1*, *HMGA2*, and *EIF3A*) also showed overwhelmingly higher expression from the primary CS compared to the subsidiary CS (**Fig. 2E, Supplemental Fig. S2B-C**). Together, these results reveal an inverse co-relation between CSH and gene expression.

**Figure 2:**
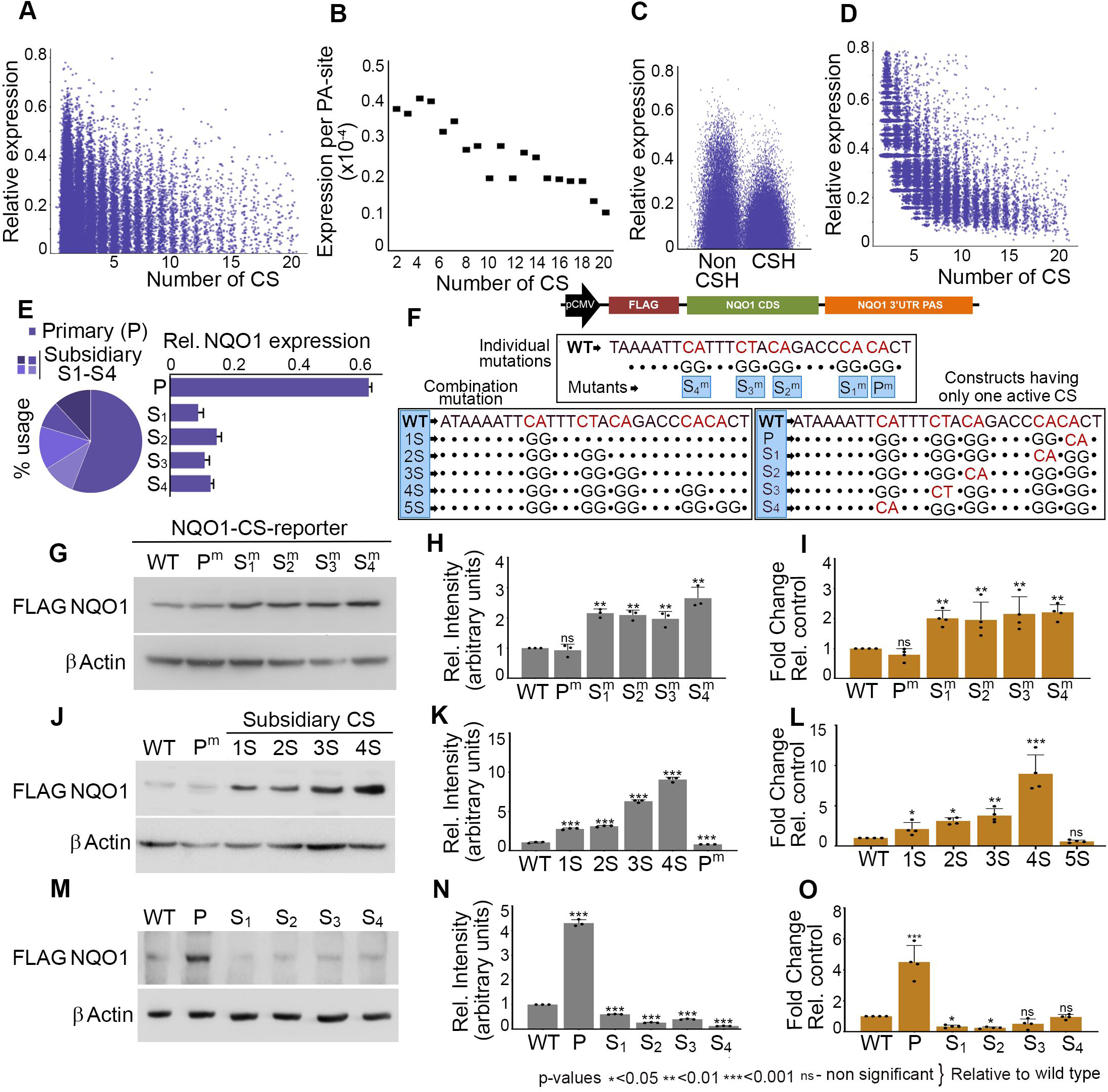
Cleavage site heterogeneity on the target PA-site regulates genes expression that exhibits an inverse correlation between the number of CS and gene expression. (*A*) Relative gene expression from each CS obtained from the total read counts per million of each CS specific transcripts of a PA-site relative to *GAPDH* level and expressed over total expression level of the particular mRNA. The relative expression is plotted against the number of PA-sites having increasing number of CS. (*B*) Average relative expression from different CS per PA-site versus the number of CS. (*C*) Relative expression as in A from PA-sites that show CSH versus that do not show CSH. (*D*) Relative expression from the primary CS as in A plotted against the number of CS being used on a particular PA-site. (*E*) Pie chart showing relative usage of different CS (primary versus subsidiary CS) of *NQO1* mRNA (left) and the bar chart of expressions from different *NQO1* CS relative to total *NQO1* expression as indicated (right). (*F*) Schematic of NQO1 terminal PA-site region showing different CS along with the different individual and combination mutations (see Supplemental Table S1). Mutant sequences are written below the WT sequence the dot (•) represents unchanged sequence. Active CS is represented in red and the mutated ones are in black. Mutant nomenclatures are highlighted in blue. (*G*) Immunoblotting of the NQO1 reporter expression from HEK293 cells using anti-FLAG-antibody with the individual mutation of the primary and different subsidiary CS as indicated. Control β-Actin is shown below. Quantification of the blot expressed relative to wild type is show in H. (*I*) qRT-PCR analysis of *NQO1* reporter expression using pair of primers (forward from the FLAG-epitope tag coding region and a reverse primer specific to *NQO1* CDS) from HEK293 cells after overexpression of wild type and different individual CS mutations as in G. (p values * <0.05, ** <0.01, *** <0.001, ns - non significant; p-values are relative to control). (*J*) Immunoblotting of the *NQO1* reporter expression from HEK293 cells using anti-FLAG-antibody with the CS combination mutations as indicated (also see Supplemental Table S1). Control β-Actin is shown below. Quantification of the blot expressed relative to wild type is show in K. (*L*) qRT-PCR analysis of *NQO1* reporter expression using a pair of primers as in I from HEK293 cells after overexpression of wild type and different combination CS mutations. Error bar represents standard error mean (SEM) of n=3 independent experiments. (p values * <0.05, ** <0.01, *** <0.001, ns - non significant; p-values are relative to control). (*M-O*) Immunoblotting followed by quantification and qRT-PCR analysis as in G-I of NQO1 reporter expression from primary and different subsidiary sites (only one CS present) in HEK293 cells. Error bar represents standard error mean (SEM) of n=3 independent experiments. (p values * <0.05, ** <0.01, *** <0.001, ns - non significant; p-values are relative to control).

### Mutational analysis of heterogenous cleavage sites confirms inverse co-relation between the cleavage site heterogeneity and protein expression

To understand the effect of CSH in gene expression, we analysed expression of *NQO1* mRNA (that encodes a key anti-oxidant protein) and specific CS. There are five major CS (one primary, P and four subsidiaries, S_1_ to S_4_) on the terminal PA-site of *NQO1* pre-mRNA. The primary CS usage was >55% and there was a corresponding higher expression from the primary CS (**Fig. 2E**). We employed a reporter mini gene construct of FLAG-tagged *NQO1* CDS under the constitutive CMV promoter and driven by the *NQO1* 3′-UTR with the terminal PA-site as described earlier(Sudheesh *et al*, 2019). We first introduced mutations in each CS (CS was mutated from CA to GG, the least efficient sequence for cleavage) (Sheets *et al*., 1990) individually by site directed mutagenesis **(Fig. 2F**, top panel). The list of mutations and sequences of mutations are listed in **Supplemental Table S1**. Reporter expression in HEK293 cells was then analysed by Western blotting using anti-FLAG antibody and qRT-PCR using a pair of primers (a forward primer from the FLAG-coding sequence and a *NQO1* CDS specific reverse primer). To our surprise, individual mutations of each subsidiary CS (S_1_^m^ to S_4_^m^) resulted in an induction of the reporter NQO1 protein and mRNA expressions (**Fig. 2G-I)**. However, there was no marked induction of NQO1 reporter expression after the mutation of the primary CS (P^m^). Next, we analysed the reporter NQO1 expression from the combination of mutations with increasing number of subsidiary CS mutations (1S, 2S, 3S and 4S) (**Fig. 2F** bottom left panel). Strikingly, we observed a gradual increase in the reporter NQO1 protein and mRNA expression with increasing number of subsidiary CS mutations (**Fig. 2J-L**). Mutation all 5 sites including the primary CS resulted in the insignificant expression of NQO1 reporter construct (**Fig. 2J-L, Supplemental Fig. S2D**). Further, to assess the expression from primary CS versus subsidiary CS, we generated CS mutants that has only one active CS where other four CS were mutated [reporter driven by only one CS, either primary CS (P), or individual subsidiary CS (S_1_-S_4_)] **(Fig. 2F** bottom right panel, **Supplemental Table S1)**. We saw highest reporter NQO1 protein and mRNA expression from the primary CS driven NQO1 reporter construct whereas the expressions from other subsidiary CS driven constructs were minimal (**Fig. 2M-O**). Consistently, there was no discernible induction of the reporter NQO1 expression with increasing number of subsidiary CS mutations after the primary CS was mutated **(Supplemental Fig. S2D-E)**. Together, these results confirm an inverse co-relation between the number of CS and the gene expression and the predominance of primary CS for the gene expression.

### Decrease in the cleavage site heterogeneity and a concomitant increase in cleavage at the primary cleavage site induces gene expression

To understand the mechanism of the inverse co-relation between the gene expression and the CSH, we analysed cellular cleavage of the reporter *NQO1* pre-mRNA construct. We saw an increase in the cleavage efficiency with each subsidiary CS mutation when the primary CS was active in agreement with the increased reporter expression (**Fig. 3A**). Moreover, there was a gradual decrease in the uncleaved pre-mRNA on increasing number of subsidiary CS mutations (**Fig. 3B**). However, in the presence of the primary CS mutation, subsidiary CS mutations did not show significant effect on the cleavage efficiency (**Supplemental Fig. S2D-E**). Moreover, we also observed highest cleavage efficiency from the primary CS compared to the other subsidiary CS (**Fig. 3C**). Concomitantly, wherever primary CS is present, predominant cleavage was observed from the primary CS irrespective of the number of subsidiary CS mutations (**Supplemental Fig. S2F**). In the absence of the primary CS, there was no predominant CS among the subsidiary CS. And there was a gradual increase in the cleavage at the primary CS with increasing number of subsidiary CS mutations (**Supplemental Fig. S2G-H**). However, when primary CS was mutated, there was no marked increase in the cleavage at any of the subsidiary CS indicating that primary CS largely drives the cleavage at the 3′-end. Together, these results reveal that increase in the primary CS usage and decrease in the number of subsidiary CS induces cleavage efficiency and subsequently the *NQO1* reporter expression.

**Figure 3:**
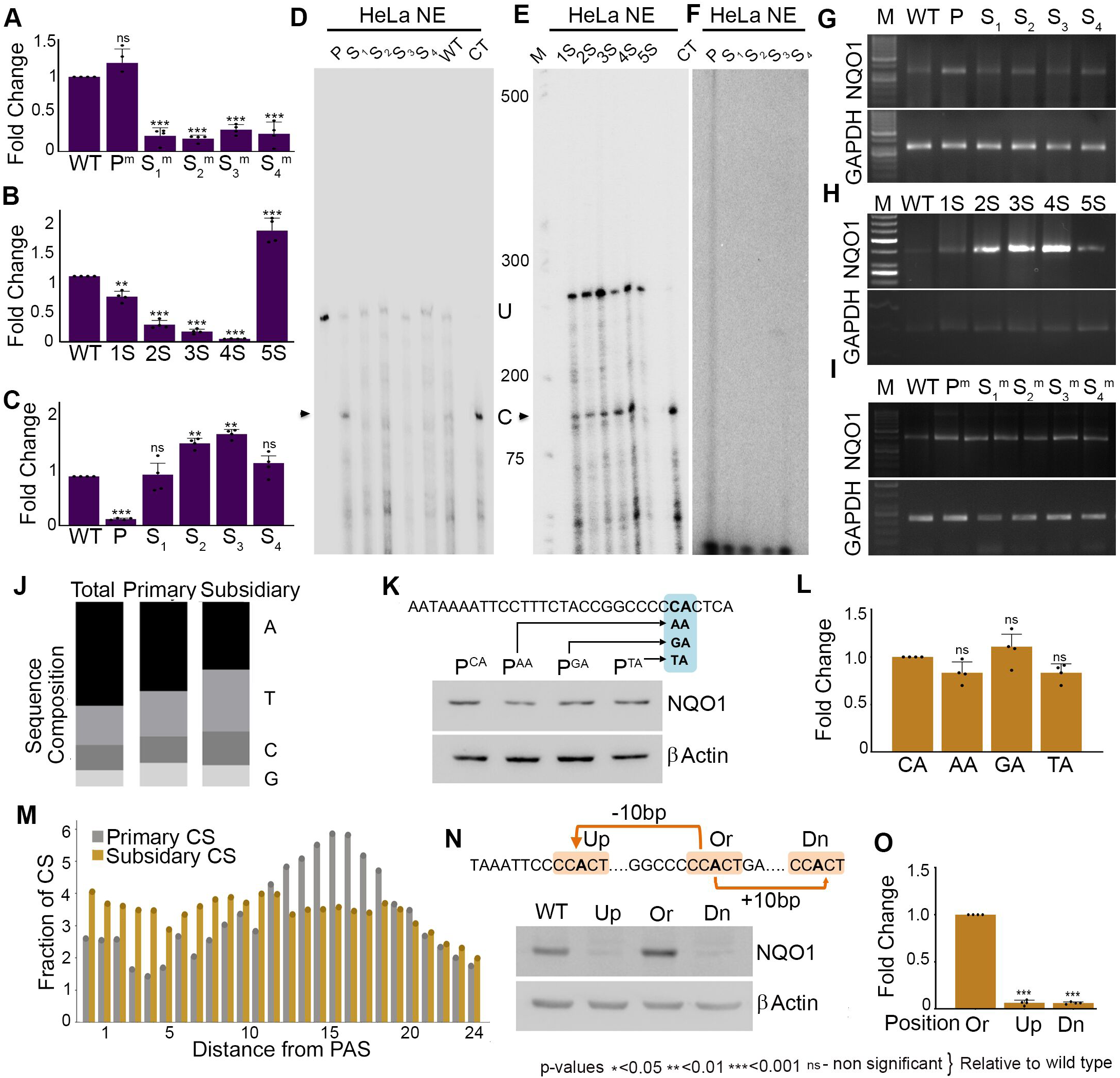
Decrease in cleavage site heterogeneity and a concomitant increase in the cleavage at the primary CS induces gene expression. (*A-C*) Measurement of in vivo cleavage efficiency of *NQO1* reporter expression from HEK293 cells after overexpression wild type and different CS mutants as in Fig. 2I, L and O. Error bar represents SEM of three independent experiments. (p values * <0.05, ** <0.01, *** <0.001, ns - non significant; p-values are relative to control). (*D*) In vitro cleavage assay of radiolabelled *NOQ1* 3′-UTR substrates (different mutants having either primary or individual subsidiary sites as in Fig. 2M) with active HeLa nuclear extracts. Control cleaved template (CT) is shown on the right. (U - uncleaved, C - cleaved fragment). (*E*) In vitro cleavage assay of radiolabelled *NOQ1* 3′-UTR substrates (wild type and different combination mutants as in Fig. 2J) with active HeLa nuclear extracts. Control cleaved template (CT) is shown on the right. (U - uncleaved, C - cleaved fragment). (*F*) In vitro polyadenylation assay of different *NQO1* cleaved templates (different mutants having either primary or individual subsidiary sites as in D) using active HeLa nuclear extract and γ-^32^P radiolabelled ATP. (*G-I*) 3′-RACE assays of NQO1 reporter expression from HEK293 cells after overexpression of wild type and different CS mutants as in A-C respectively. (*J*) Sequence composition and frequency of occurrence of the terminal CS nucleotide (A, T, G, C) of primary and subsidiary CS genome-wide. (*K*) Immunoblotting of the NQO1 reporter expression from HEK293 cells after overexpression of the CS sequence mutants (P^CA^, P^AA^, P^GA^, P^TA^) in the HEK293 cell. (L) qRT-PCR analysis of *NQO1* reporter expression after overexpression of the CS sequence mutants (P^CA^, P^AA^, P^GA^, P^TA^) in the HEK293 cell. Error bar represents SEM of three independent experiments. (p values * <0.05, ** <0.01, *** <0.001, ns - non significant; p-values are relative to control). (*M*) Distance of the occurrence of the primary versus subsidiary CS relative to the PA-signal on each PA-site. The primary and subsidiary CS are indicated. (*N*) Immunoblotting of the NQO1 reporter expression (wild type and position mutations of the primary CS) and control β-Actin from HEK293 cells after overexpression of the respective mutants. The original position of the primary CS (Ori) was moved 10 nucleotides upstream (Up) or downstream (Dn) as indicated. Quantification of the blots with mutants relative to the original is shown in Supplemental Fig. 3F. (*O*) qRT-PCR analysis of *NQO1* reporter expression (with forward primer from the FLAG-epitope tag coding region and a reverse primer specific to *NQO1* CDS) from HEK293 cells after overexpression of wild type and position mutations as in L. Error bar represents SEM of three independent experiments. (p values * <0.05, ** <0.01, *** <0.001, ns - non significant; p-values are relative to control).

To confirm this finding, we then carried out an in vitro cleavage assay using 5′-capped *NQO1* 3′-UTR RNA encompassing the sequences required for cleavage and polyadenylation (∼120 nucleotides upstream and downstream of the CS of the terminal PA-site) and HeLa nuclear extract as reported earlier(Laishram & Anderson, 2010; Sudheesh *et al*., 2019). Consistently, we observed highest cleavage efficiency from the primary CS containing *NQO1* 3′-UTR RNA, whereas the cleavage from the subsidiary CS were negligible in vitro (**Fig. 3D, Supplemental Fig. S3A**). Akin to that in cellular assays, we also saw a gradual increase in the cleavage efficiency with decreasing number of subsidiary CS (**Fig. 3E, Supplemental Fig. S3B**). Since, cleavage is coupled with polyadenylation, we also assessed the downstream polyadenylation efficiency using in vitro transcribed cleaved templates with each CS. We found the highest polyadenylation efficiency from the templates cleaved at the primary CS than the other subsidiary CS that showed negligible polyadenylation efficiency in vitro (**Fig. 3F, Supplemental Fig. S3C**). To confirm this, we then performed 3′-RACE to measure the 3′-end formation with wild type and mutant *NQO1* reporter constructs (**Fig. 3G-I**). We saw highest polyadenylation efficiency of the primary CS driven *NQO1* reporter transcript with a gradual enhancement of polyadenylation with each subsidiary CS mutation (**Fig. 3G-I**). However, we did not see any significant difference in the length of the PA-tail addition on the primary and subsidiary CS specific transcripts **(Supplemental Fig. S3D)**. These results confirm that primary CS exhibits highest cleavage and polyadenylation efficiency that drives gene expression.

### Position of the CS downstream of the PA-signal but not the sequence composition determines the primary CS

Next, we investigated what determines the primary CS, we looked into two key determinants: sequence composition and the nucleotide position from the PA-signal. Earlier studies established the predominant nucleotide at the CS (A>T>C>>G) and the dinucleotide (CA>AA>TA>>GA) in the order(Chen *et al*., 1995). We first analysed the terminal nucleotide sequence of primary and subsidiary CS genome wide (**Fig. 3J**). We observed a similar occurrence of the terminal nucleotide and the dinucleotide at the CS as shown earlier for general CS (terminal nucleotide: A>T>C>G, dinucleotide CA>AA>TA>GA) in both primary and subsidiary CS (**Fig. 3J, Supplemental Fig. S3E**). This indicates that sequence composition is unlikely to determine the primary versus subsidiary CS. Consistently, when we mutated the primary CS dinucleotide (-CA-) to other dinucleotide combinations (CA to -TA-, -GA-, and - AA-) , there was no significant difference in the reporter NQO1 protein and mRNA expressions between the wild type sequence and the mutant sequences of the primary CS (**Fig. 3K-L**). Likewise, there was no change in the 3′-end formation between the wild type and mutant sequences in the reporter analysis (**Supplemental Fig. S3F-G**). We then analysed the position of occurrence of the primary CS and subsidiary CS for all the PA-sites genome wide. We observed primary CS largely clustered between positions 12 to 18 nucleotides downstream of the PA-signal whereas subsidiary CS did not show any specific clustering and were randomly distributed (**Fig. 3M**). In majority of the PA-sites, the primary CS was located between 14 to 18 nucleotides (a median value of ∼15.6 nucleotides) downstream of the PA-signal (**Fig. 3M**) showing a positional bias for the occurrence of primary CS relative to the PA-signal. To confirm this positional bias of the primary CS, we employed *NQO1* reporter construct with only the primary CS at the 3′-UTR. We introduced mutations to alter the position of the primary CS by moving the CS region (a sequence of 5 nucleotides, two nucleotides flanking the CS, CC*A*CT where the CS is italicised) to different position from the original position (16 nucleotides downstream of the PA-signal). We moved the CS sequence to 10 nucleotides upstream in one construct and 10 nucleotides downstream of the original position in another construct (**Fig. 3N**) (**Supplemental Table S1**). Surprisingly, we saw a drastic reduction in the *NQO1* reporter expression when the position of CS was changed to upstream and downstream positions relative to the original position (**Fig. 3N-O, Supplemental Fig. S3H**). There was >5-fold difference in the expression between the reporter expression from original primary CS position and the new changed positions. Consistently, we saw an increased accumulation of uncleaved pre-mRNA and compromised 3′-end formation when the position of the primary CS was changed from its original position (**Supplemental Fig. S3I-J**). Together, these results demonstrate that the position of the CS relative to the PA-signal and not the sequence composition determines the primary CS of a PA-site.

### Reduced cleavage heterogeneity is directly associated with oxidative stress response gene expression

To understand the biological relevance of CSH, we first performed enrichment analysis of genes that shows CSH in different cellular pathways in silico (Metascape). Interestingly, we observed gene enrichment in pathways directly or indirectly involved in cellular response to different stresses that incudes oxidative stress, ER stress, chemical induced stress, mitochondrial stress, nutrient stress, unfolded protein response etc. (**Supplemental Fig. S4A**) (**Supplemental Dataset 2**). Next, we checked important genes related with oxidative stress response (>800 genes detected in earlier studies) (**Supplemental Table S2**) (Fabregat *et al*, 2018). Strikingly, >90% of the PA-sites of these genes showed heterogeneity **(Fig. 4A)**. The median number of CS of the oxidative stress response genes were 4.5 compared to the 1.5 of the randomly selected equivalent number of non-stress response genes (**Fig. 4B**). 20 key oxidative stress response genes along with their number of CS is shown in **Supplemental Fig. S4B**. Concomitantly, mRNA isoforms of stress response genes showed lower relative expression than that of non-stress response genes under normal condition (**Fig. 4C**) consistent with the higher CSH. To understand the implication of CSH in oxidative stress response gene expression, we then analysed gene expression versus CSH after treatment with two stressors that induces anti-oxidant response - hydrogen peroxide (H_2_O_2_) and sodium arsenite (NaAsO_2_) from earlier studies (Mullani N, 2021; Zheng *et al*, 2018)**(Fig. 4D-G)**. There was a global induction of oxidative stress response genes under the two stress conditions. Consistently, we observed a global decrease in the CSH and an increase in the primary CS usage on both H_2_O_2_ and NaAsO_2_ treatment **(Fig. 4D-G)**. Under H_2_O_2_ treatment, the median number of CS was reduced from 6.8 to 1.5 whereas the primary CS usage was increased from 50% average usage to >80% average usage after treatment with H_2_O_2_ **(Fig. 4D-E)**. Likewise, under NaAsO_2_ treatment, there was a similar decrease in the CSH from 4.5 median CS to 2.2 median CS, whereas the primary site usage was increased from 55% to 75% (**Fig. 4F-G**) indicating that CSH and primary CS usage is directly linked with the oxidative stress response gene expression. Concomitantly, when we look at the individual stress response genes, we saw an equal reduction in the CSH and increase in the primary CS usage on stress (NaAsO_2_) treatment (**Supplemental Fig. S4C**). Together, these results reveal that induction of anti-oxidant gene expression is associated with a decrease in the CSH and an increase in the primary CS usage.

**Figure 4:**
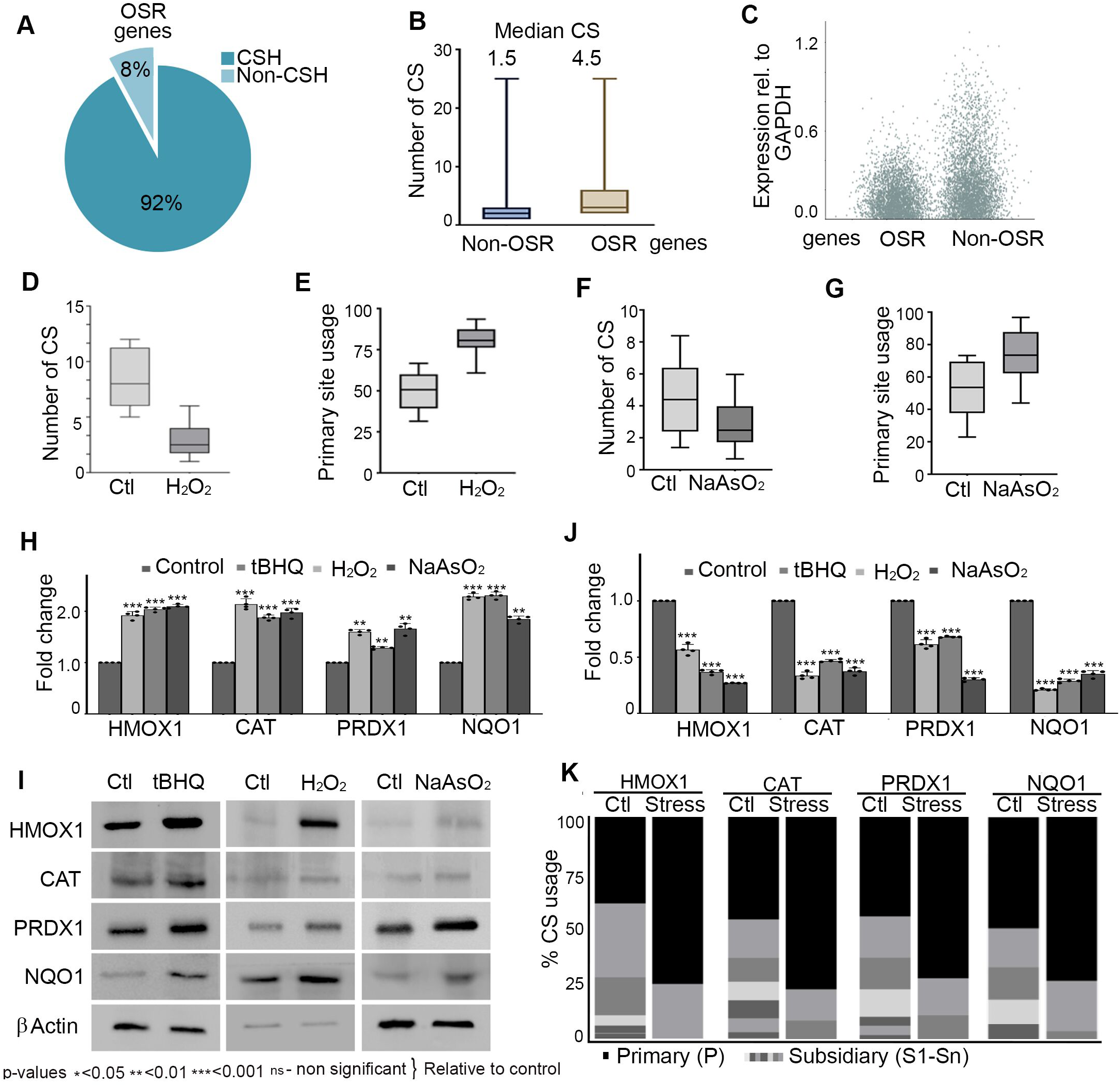
Reduced cleavage heterogeneity regulates increased anti-oxidant gene expression during oxidative stress response. (*A*) Pie chart showing fraction of mRNAs that shows CSH and non-CSH among the genes encoding oxidative stress response proteins. (OSR - Oxidative stress response genes) (*B*) Box plot of frequency of CSH of PA-sites from oxidative stress related and random genes as in B. Median CS of PA-sites is indicated. (OSR - Oxidative stress response genes) (*C*) Relative expression of CS as in Fig. 2C from PA-sites of control and stress response genes. (OSR – Oxidative stress response genes). (*D-E*) Box plot showing reduction in the CSH and increase in % usage of the primary CS of stress response genes in presence of H_2_O_2_ induction. Data was analysed from earlier sequencing data (GSE138290)(Mullani N, 2021). (*F-G*) Box plot Box plot showing reduction in the CSH and increase % usage as in E-F but in presence of NaAsO_2_ treatment. Data was analysed from earlier sequencing data (GSE101851)(Zheng *et al*., 2018). (*H*) qRT-PCR of stress response mRNAs *HMOX1, CAT, PRDX1* and *NQO1* in HEK293 cell in the presence and absence of oxidative stress induction in HEK293 cells with different stressors (tBHQ, H_2_O_2_, NaAsO_2_ as indicated). (p values * <0.05, ** <0.01, *** <0.001, ns - non significant; p-values are relative to control). (*I*) Immunoblotting of the HMOX1, CAT, PRDX1, NQO1 and control β-Actin under the conditions as indicated. (*J*) Measurement of in vivo cleavage efficiency of stress response mRNAs from HEK293 cells as in I. (p values * <0.05, ** <0.01, *** <0.001, ns - non significant; p-values are relative to control). (*K*) Stacked column plot of % usage of primary versus subsidiary CS of stress response mRNAs *HMOX1, CAT, PRDX1, NQO1* in the presence and absence of oxidative stress induction (similar results were obtained for all stressors tBHQ, H_2_O_2_, NaAsO_2_). Primary CS is indicated in black and different subsidiary CS are indicated in different shades of grey.

### Cleavage heterogeneity regulates expression of oxidative stress response proteins under oxidative stress

Next, to further assess the role of CSH in antioxidant gene expression, we employed three stressors H_2_O_2_, NaAsO_2_, and the oxidative stress agonist tert-butyl hydroquinone (tBHQ) and induced anti-oxidant response in HEK293 cells. We then analysed the expression and CS usage of four key oxidative stress response genes (*HMOX1*, *CAT*, *PRDX1*, and *NQO1*). We consistently observed induced protein and mRNA expressions of the four-stress response regulator assessed on treatment with the three stressors (**Fig. 4H-I**). Analysis of uncleaved pre-mRNA also showed increased cleavage efficiency of the selected four stress response pre-mRNAs on treatment with the three different stressors (**Fig. 4J**). In agreement with this, we also saw a reduced CSH of the four genes on stress induction (**Fig. 4K**). The number of CS was 6 for *HMOX1*, 8 for *CAT* and *PRDX1*, and 5 for *NQO1* respectively under normal condition that was reduced to 2 CS for *HMOX1* and *NQO1*, and 3 CS for *CAT* and *PRDX1* under all the three stress conditions (**Fig. 4K**). Concomitantly, there was also an increase in the primary CS usage from ∼45-50% under normal condition to >70% under all stress conditions (**Fig. 4K, Supplemental Fig. S5A**) confirming the association of CSH with anti-oxidant gene expression.

Next, we used NQO1 reporter mini gene construct in the presence and absence of three stressors (H_2_O_2_, NaAsO_2_, and tBHQ). First, we validated the induced *NQO1* reporter expression by Western blot and qRT-PCR analysis along with in vivo cleavage after treatment with the three stressors (**Fig. 5A, Supplemental Fig. S5B-C**). All three stressors showed stimulations of *NQO1* reporter expression and cleavage efficiency. Moreover, we saw a similar reduction in the CSH and increase in the primary CS usage on treatment with the three stressors (**Supplemental Fig. S5D**). To confirm the role of CSH in the stress response gene expression, we employed the wild type reporter construct that shows CSH and the mutant construct lacks the heterogeneity (having only the primary CS and other subsidiary sites were mutated). Strikingly, we observed no induction of the reporter *NQO1* expression from the mutant construct that lacks CSH whereas there was induced expression from the wild type construct on stress treatment **(Fig. 5B)**. Similarly, *NQO1* expressions from the reporter construct that has only one individual subsidiary CS was not stimulated on stress induction (**Supplemental Fig. S5E, F**) indicating that CSH is involved in the anti-oxidant gene expression. Consistently, expression from the primary CS was highest compared to the subsidiary CS under all the three stress conditions **(Supplemental Fig. S5G)**. To further assess the role of CSH, we checked the reporter expression using *NQO1* constructs with increasing number of subsidiary CS mutations. We observed a gradual increase in the induced *NQO1* protein expression with increasing number of subsidiary site mutations on stress treatment (**Fig. 5C-D, Supplemental Fig. S5H-I**). Likewise, there was an increase in the mRNA level and cleavage efficiency from the reporter expression with the subsidiary CS mutations (**Fig. 5E-F**). However, the highest difference in the expression level between control and stress treatment was seen from the wild type reporter construct having 4 subsidiary CS, and least when all subsidiary CS were mutated (**Supplemental Fig. S5J-K**). Together, these results showed that stress response gene expression involves both reduction in the CSH along with increase in the primary CS usage. And that when only primary CS is present, the expression becomes constitutive.

**Figure 5:**
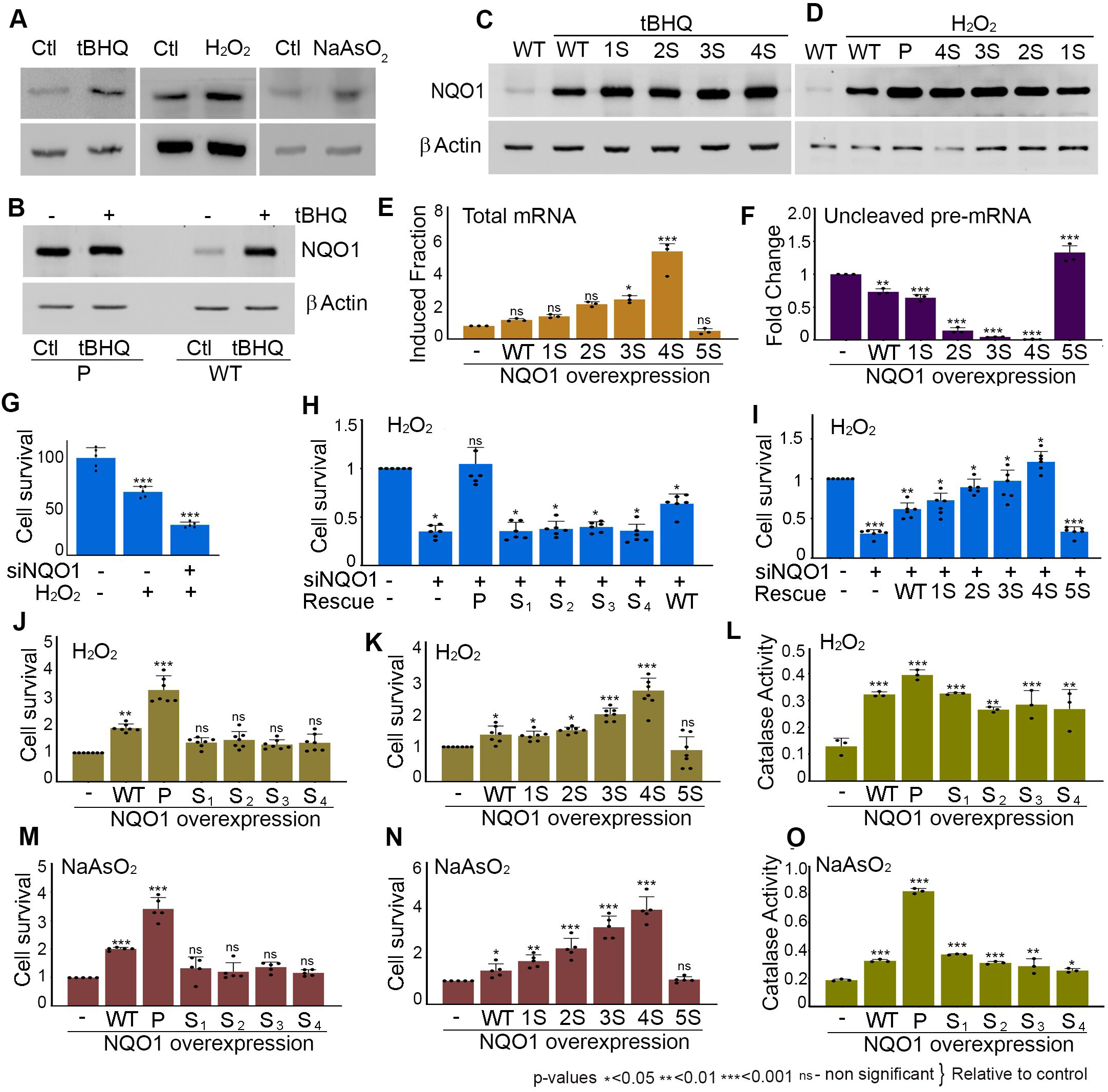
Cleavage heterogeneity regulates cellular oxidative stress response. (*A*) Immunoblotting of the wild type FLAG-NQO1 reporter expression and control β-Actin in HEK293 cell after induction of oxidative stress using tBHQ, H_2_O_2_, and NaAsO_2_. (*B*) Immunoblotting of wild type and mutant *NQO1* reporter (having only the primary CS) expression and control β-Actin in HEK293 cell after treatment with tBHQ. (*C-D*) Immunoblotting of the NQO1 reporter expression and control β-Actin after overexpression of reporter constructs with increasing subsidiary CS mutations as indicated in HEK293 cell in the presence of induction of oxidative stress using tBHQ and H_2_O_2_ respectively. Blot of untreated control is also indicated. (*E*) qRT-PCR of induced *NQO1* reporter expression relative to uninduced expression from HEK293 cells with increasing subsidiary site mutations upon H_2_O_2_ treatment. Error bar represents standard error mean (SEM) of n=3 independent experiments. (p values * <0.05, ** <0.01, *** <0.001, ns - non significant; p-values are relative to control). (*F*) In vivo cleavage assay of the *NQO1* reporter expression from HEK293 cells as in E. Error bar represents standard error mean (SEM) of n=3 independent experiments. (p values * <0.05, ** <0.01, *** <0.001, ns - non significant; p-values are relative to control). (*G*) MTT assay to assess cell viability in HEK293 cells after H_2_O_2_ treatment in the presence and absence of knockdown of *NQO1*. (*H*) MTT assay as in G after knockdown of *NQO1* and rescue with wild type and different *NQO1* CS mutants having only primary or individual subsidiary sites as indicated. (*I*) MTT assay as in H but in rescue with wild type and *NQO1* CS mutants with increasing subsidiary site mutations as indicated. (*J*) MTT assay in HEK293 cells after overexpression of wild type and different *NQO1* CS mutants having only primary or individual subsidiary sites as indicated after induction of oxidative stress by H_2_O_2_. (*K*) MTT assay in HEK293 cells after overexpression of wild type and different *NQO1* CS mutants with increasing subsidiary CS mutations as indicated after induction of oxidative stress by H_2_O_2_. (*L*) Catalase assay HEK293 cells under the similar conditions as in J. (*M-N*) MTT assay as in J-K respectively but after induction of oxidative stress with NaAsO_2_ treatment. (*O*) Catalase assay as in L after induction of oxidative stress with NaAsO_2_. For all MTT and catalase assays, error bar represents standard error mean (SEM) of n=3 independent experiments (p values * <0.05, ** <0.01, *** <0.001, ns - non significant; p-values are relative to control).

### Cleavage site heterogeneity versus primary cleavage site usage regulates cellular tolerance to oxidative stress induction

Since NQO1 is a key antioxidant response protein, we tested the effect of *NQO1* expression from reduced CSH in cellular oxidative stress tolerance in HEK293 cells using two stressors H_2_O_2_ and NaAsO_2_. We first knocked down *NQO1* in HEK293 cells and treated cells with H_2_O_2_ and rescued with wild type and *NQO1* CS mutant constructs that are insensitive to the *NQO1* siRNA employed. Treatment with H_2_O_2_ resulted in a significant cellular sensitivity with around 40% cell death in control cells in MTT assay (**Fig. 5G**). NQO1 knockdown resulted in a further sensitivity with an overall >60% cell death compared to control cells on the H_2_O_2_ treatment (**Fig. 5G**). Intriguingly, *NQO1* expression from the primary CS but not the other subsidiary CS rescued the cell death from the knockdown of *NQO1* on H_2_O_2_ treatment (**Fig. 5H**). Expression from the wild type *NQO1* construct also increased the stress tolerance, yet the highest tolerance to cellular stress was obtained from the primary CS (**Fig. 5H**). Similarly, rescue of the H_2_O_2_ sensitivity was gradually increased with increasing number of subsidiary CS mutations on stress treatment (**Fig. 5I**) in agreement with the increased *NQO1* expression. Further, we overexpressed NQO1 from different ectopic constructs with individual and combination of subsidiary CS mutations and assessed the cellular tolerance to H_2_O_2_ treatment. Akin to earlier results, ectopic overexpression of *NQO1* from the wild type construct increased cellular tolerance to H_2_O_2_ stress with a highest tolerance from *NQO1* expression from the primary CS. *NQO1* expression from subsidiary CS did not impart marked effect on the tolerance to H_2_O_2_ stress (**Fig. 5J**). We also saw a gradual increase in the cellular H_2_O_2_ tolerance with increasing number of subsidiary CS mutations (**Fig. 5K**). Similar cell sensitivity was also scored by trypan blues staining after H_2_O_2_ treatment in the presence and absence of ectopic *NQO1* expression from wild type and mutant CS constructs. We observed similar results in trypan blue staining with a loss of tolerance from the primary CS mutation and a gradual increase in the tolerance with decreasing CSH (**Supplemental Fig. S6A-B**). Moreover, assessment of the catalase activity after H_2_O_2_ treatment also showed higher catalase activity on *NQO1* expression from the primary CS having no CSH compared to the control cells (**Fig. 5L**) revealing the role of CSH in the cellular anti-oxidant response.

Concomitantly, stress tolerance was tested after NaAsO_2_ treatment in the presence of ectopic NQO1 expression from individual CS and combination CS mutations that showed a gradual decrease in the CSH. We found a similar increase in the stress tolerance with *NQO1* expression from the primary CS driven construct than the other subsidiary CS driven constructs (**Fig. 5M**). There was >1.8-fold increase in the cell survival in MTT assay from *NQO1* expression from the primary CS relative to the control cells. Moreover, there was a gradual increase in the tolerance with increasing number of mutations of the subsidiary CS (**Fig. 5N**) as seen in H_2_O_2_ treatment. Similar results were obtained in the trypan blue staining after NaAsO_2_ treatment as in the case of MTT assay. There was >2.5-fold increase in the cell survival on NQO1 expression compared to the control cells with no ectopic expression. The tolerance was lost on the primary CS mutation, whereas, the individual subsidiary CS mutation had only modest effect on the sensitivity (**Supplemental Fig. S6C**). Moreover, there was also a gradual increase in the NaAsO_2_ tolerance with NQO1 expression with decreasing CSH (**Supplemental Fig. S6D**). Furthermore, we also measured the catalase activity after NaAsO_2_ treatment and we observed highest catalase activity in the presence of NQO1 expression from the primary CS than that of wild type or other subsidiary CS **(Fig. 5O**). Together, these results demonstrate a correlation between CSH and cellular tolerance to oxidative stress and that the anti-oxidant response involves both decrease in the CSH and increase in the primary CS usage.

### Increased affinity of cleavage complex assembly stimulates fidelity of cleavage and reduces cleavage heterogeneity upon oxidative stress induction

Earlier studies showed that oxidative stress response gene expression is regulated by the non-canonical PAP, Star-PAP and not by the canonical PAPα and that Star-PAP activity is induced during oxidative stress through phosphorylation (Gonzales *et al*, 2008; Mellman *et al*., 2008). We also confirmed the expression of key oxidative stress response regulators (*NQO1*, *HMOX1*, *PRDX1*, *CAT*) by Western and qRT-PCR. We observed loss of both protein and mRNA expressions on Star-PAP knockdown but not on PAPα knockdown (**Supplemental Fig. S6E-F**). To understand the mechanism of CSH and how CSH is reduced and the primary CS usage is stimulated during oxidative stress, we analysed the Star-PAP cleavage complex assembly in the presence and absence of oxidative stress agonist, tBHQ treatment. We first immunoprecipitated Star-PAP and analysed the association of key CPA component (CPSF subunits 73, 30, 100, and 160). Intriguingly, we observed induced association of Star-PAP with CPSF components CPSF30, CPSF73 and CPSF100 suggesting an increased affinity of Star-PAP CPA complex under oxidative stress (**Fig. 6A**,). Since Star-PAP binds target mRNAs and helps recruit cleavage components CPSF73 and CPSF160 through direct interaction (Kandala *et al*, 2016; Laishram & Anderson, 2010), we assessed the implication of increased affinity of the complex assembly on the target mRNA association by RNA immunoprecipitation (RIP) using Star-PAP and CPSF73 pulldown. We found heightened association of both Star-PAP and CPSF73 on mRNAs encoding key anti-oxidant response protein (*NQO1, HMOX1, PRDX1*) upon oxidative stress induction **(Fig. 6B)**. There was 3- to 5-fold increase in the association of Star-PAP and CPSF73 on the target mRNAs on tBHQ treatment (**Supplemental Fig. S6G-H**) revealing that oxidative stress induces the affinity of the Star-PAP cleavage complex assembly on mRNA target 3′-UTRs. To understand the link between the increased affinity of CPA complex assembly with CSH, we employed the NQO1 reporter construct with wild type and CSH mutant (having only the primary CS that do not show CSH) and analysed the association of CPSF73 and Star-PAP by RIP experiment. Consistently, we saw increased association of Star-PAP and CPSF73 with the *NQO1* with the wild type 3′-UTR (that shows CSH) on tBHQ treatment **(Fig. 6B, Supplemental Fig. S6G-H)**. However, in the presence of CSH mutant, there was no further induction in the association of *NQO1* 3′-UTR with CPSF73 and/or Star-PAP in the reporter RNA indicating that CSH is linked with the increased affinity of the CPA component assembly with the target mRNA.

**Figure 6:**
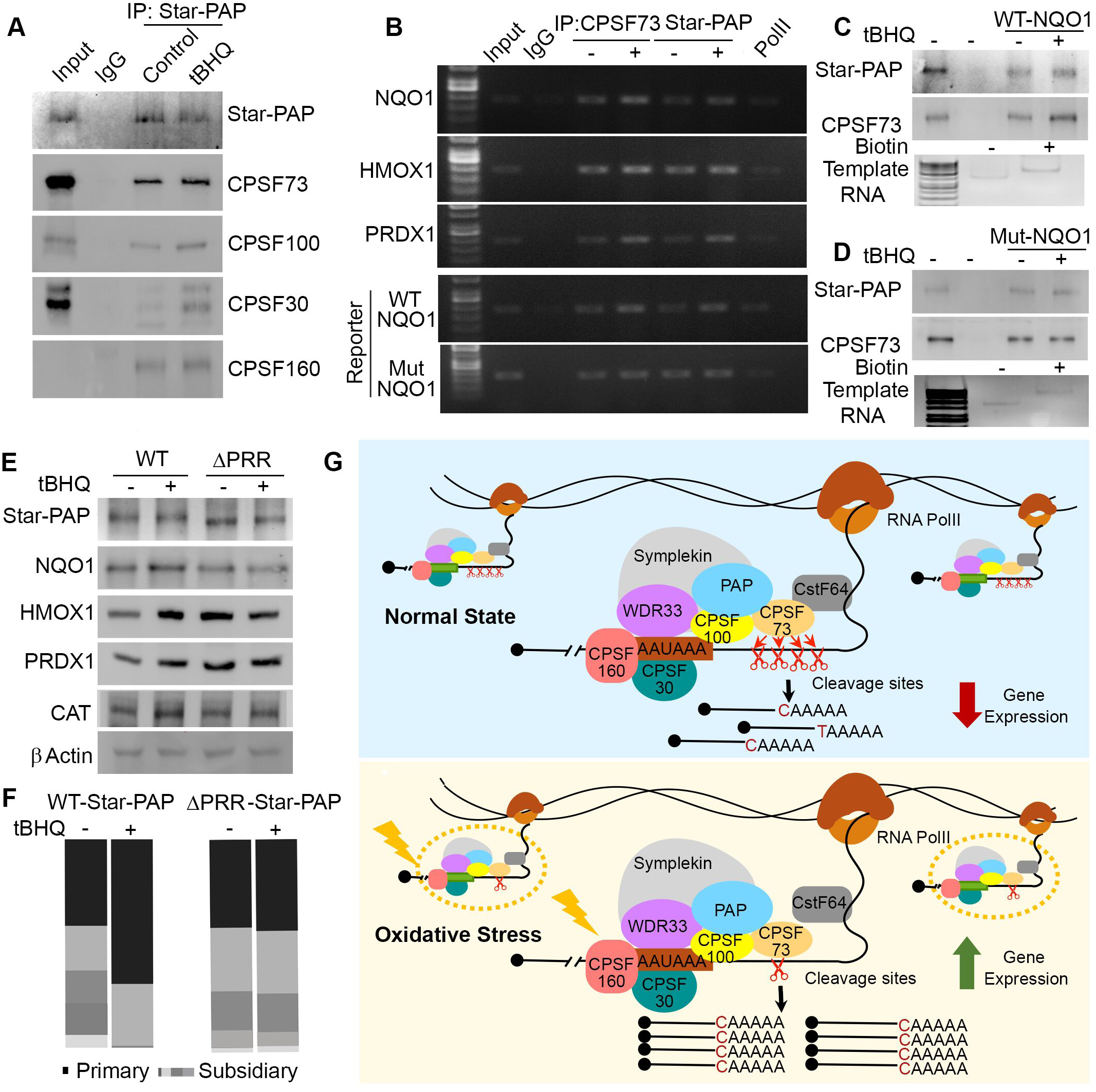
Increased affinity of cleavage complex assembly on oxidative stress stimulates fidelity of cleavage at the primary site reducing the CSH. (*A*) Immunoprecipitation of Star-PAP from HEK293 cells in the presence and absence of oxidative stress induction by tBHQ followed by Western blot analysis of Star-PAP, CPSF73, CPSF100, CPSF30 and CPSF160 as indicated. Input is equivalent of 10% of the total protein of the IP lysate used. (*B*) RNA immunoprecipitation (RIP) analysis of Star-PAP and CPSF73 along with control RNA pol II from HEK293 cells in the presence and absence of tBHQ treatment and the analysis of association with oxidative stress response genes (*NQO1, HMOX1, PRDX1*) and also association with reporter WT and mutant *NQO1* RNA is indicated. Input is equivalent of 10% of the total protein of the IP lysate used. (*C*) Streptavidin pulldown of biotin labelled wild type *NQO1* from cell extracts followed by Western analysis of Star-PAP and CPSF73in the presence and absence of tBHQ induction. Biotinylated and non-biotinylated RNA templates are shown below. (*D*) Streptavidin pulldown of biotin labelled mutant *NQO1* from cell extracts followed by Western analysis of Star-PAP and CPSF73 in the presence and absence of tBHQ induction. Biotinylated and non-biotinylated RNA templates are shown below. (*E*) Immunoblotting analysis of oxidative stress response proteins (NQO1, HMOX1, PRDX1) from HEK293 cells in the presence and absence of tBHQ treatment after Star-PAP knockdown and rescue with wild type and PRR deletion (ΔPRR) Star-PAP. (*F*) Stacked column plot of different CS usage in HEK293 cells of *NQO1* terminal PA-site under the same condition as in F. (*G*) Schematics of cleavage site heterogeneity and its regulation in oxidative stress response.

To confirm this, we prepared biotinylated *NQO1* 3′-UTR RNA templates with wild type and the CSH mutant 3′-UTR (having only primary CS, other subsidiary sites were mutated) by in vitro transcription. We then incubated both wild type and CSH mutant biotinylated 3′-UTR RNAs with active nuclear extracts and then pulled down the associated proteins using streptavidin beads. We then analysed the associated cleavage factors by Western blotting. We consistently observed increased association of CPSF73 and Star-PAP with the wild type RNA on tBHQ treatment (**Fig. 6C**). However, there was no further increase in the association of CPSF73 or Star-PAP with the mutant *NQO1*-UTR on tBHQ treatment (**Fig. 6D**) confirming a direct link between CPA complex assembly and the CSH. Together, these results suggest that increase affinity of the CPA complex assembly increases the fidelity of cleavage at the primary CS thus reducing CSH. To test this, we first looked for Star-PAP deletion mutations that will compromise stimulated association of the CPSF components with Star-PAP. We employed deletion mutations of several Star-PAP domains that were earlier linked with oxidative stress covering most of the Star-PAP CDS region [ZF-RRM deletion (ΔZF-RRM), proline rich region in the PAP domain deletion (ΔPRR), and C-terminal deletion (ΔCTD)]. Strikingly, we observed a loss of induced association of CPSF73, CPSF30 and CPSF100 with Star-PAP on tBHQ treatment on PRR deletion (**Supplemental Fig. S6I**). ΔZF-RRM or ΔCTD Star-PAP showed similar induction in the association with CPSF components as in the case of wild type Star-PAP. Star-PAP PRR region and its phosphorylation has established role in the oxidative stress response (Gonzales *et al*., 2008; Mellman *et al*., 2008). Star-PAP ΔPRR can now act as Star-PAP mutant that is compromised for induced CPA complex assembly under oxidative stress condition. Therefore, we then tested the oxidative stress response gene expression and CSH with the wild type and ΔPRR mutant Star-PAP in HEK293 cells after endogenous depletion of Star-PAP. Strikingly, ΔPRR Star-PAP failed to show induced expression of oxidative stress response proteins (NQO1, HMOX1, PRDX1, and CAT) and respective mRNA expression on tBHQ treatment while wild type expression showed induced levels on stress treatment (**Fig. 6E, Supplemental Fig. 6J**). Strikingly, assessment of the CSH of reporter *NQO1* 3′-UTR showed no change in the CSH in the presence of ΔPRR mutation whereas the wild type Star-PAP showed reduction in the CSH (**Fig. 6F**). There was ∼60% decrease in the heterogeneity in the presence of wild type Star-PAP on stress induction, there was no significant reduction in the CSH in the presence of ΔPRR mutant Star-PAP. Moreover, while there is ∼40% increase in the reporter *NQO1* 3′-UTR primary CS usage in the presence of wild type Star-PAP, there was no increase in the primary usage in the presence of ΔPRR mutant Star-PAP (**Fig. 6F**). Together, these results reveal that oxidative stress induction stimulates the affinity of CPA assembly thereby increasing the fidelity of cleavage at the primary CS and reducing the CSH upon oxidative stress treatment.

## DISCUSSION

3′-end polyadenylation involves two distinct yet coupled steps - endonucleolytic cleavage and polyadenylation (Chan *et al*, 2011; Sun *et al*, 2020; Wahle & Keller, 1996). Cleavage reaction influences the efficiency of mature mRNA template generation for the downstream polyadenylation. Consequentially, mutation in the CS compromises the overall 3′-end processing reaction, and such mutations are known in human genetic diseases(Bishop *et al*., 1988; Sheets *et al*., 1990). Yet, cleavage reaction is considered stochastic occurring downstream of the PA-signal that can result in the heterogeneity in CS(Chen *et al*., 1995; Pauws *et al*., 2001; Stroup & Ji, 2023). Contrary to this, we demonstrated that cleavage reaction is tightly regulated through CSH to control gene expression and that the heterogeneity is not a result of imprecision or random cleavage events. In the CSH, cleavage occurs predominantly at a primary CS that is followed by cleavages at different subsidiary sites. We demonstrated an inverse relationship between the number of CS and the gene expression that regulates cellular anti-oxidant gene expression. During the anti-oxidant response, there is a decrease in the cleavage at the subsidiary CS while increase in the usage of the primary CS that stimulates the expression of anti-oxidant response proteins. This mechanism is shown in **Fig. 6G**. This mechanism accounts for almost all mRNAs related to stress response gene expression inducing the cleavage and polyadenylation efficiency. Therefore, it is likely that the imprecision of cleavage reaction normally occurs in the cell and that the precision is improved under oxidative stress leading to reduced CSH and induced gene expression.

Star-PAP is the primarily PAP that is involved in the regulation of oxidative stress response gene expression(Gonzales *et al*., 2008; Mellman *et al*., 2008). Star-PAP polyadenylation activity is dramatically stimulated during antioxidant response(Mellman *et al*., 2008). Therefore, the increase in the polyadenylation activity will complement the increased cleavage efficiency thus yielding a stimulated and rapid production of mature mRNAs encoding anti-oxidant response proteins. Moreover, Star-PAP has direct role in the cleavage step that recruits the endonuclease CPSF73 in addition to acting as a structural protein to assemble the cleavage complex on target mRNAs (Kandala *et al*., 2016; Laishram & Anderson, 2010). Our reporter analysis reveals that the mechanism of CSH is distinct from other transcriptional mechanism of stress response gene expression(de Nadal *et al*, 2011; Tonelli *et al*, 2018). CSH-mediated mechanism at the 3′-end is likely to function in concert (or in collaboration) with the transcriptional mechanism at the 5′-end preventing futile cycle of transcriptional stimulation and inducing the overall gene expression. Hence, CSH-mediated gene regulation will have ramifications in the disease pathogenesis such as cardiovascular, cancer, inflammation, pathogenesis, neurodegeneration, aging, or diabetes where anti-oxidant response is critical(Cheng *et al*, 2022; Hajam *et al*, 2022; Jelic *et al*, 2021; Raut & Khullar, 2023; Saleem *et al*, 2020).

Next is the mechanism of CSH and its regulation. We have answered this question in the case of oxidative stress response gene expression. Anti-oxidant response involves both reduction in the CSH and increase in the primary CS usage. We have shown affinity of CPA complex is induced under oxidative stress that increases the strength of CPA complex assembly. We propose that increase in the strength of the CPA complex assembly induces the fidelity of cleavage at the primary CS thus reducing the suboptimal heterogenous cleavages. Our study directly links the affinity of CPA complex assembly with CSH and shows tight regulation of CSH where cleavage occurs largely at the primary CS followed by a number of random suboptimal cleavages at nearby sites. Earlier studies had shown lower efficiency of cleavage with compromised cis-elements at the 3′-UTR (Abbass *et al*, 1997; Montell C, 1983). In the light of our new finding, it is likely because of lower affinity/strength of CPA assembly resulting in increased CSH reducing the cleavage efficiency. Moreover, a recent study also suggested potential relationship between the strength of CPA complex assembly with CSH, where decrease in the strength of cleavage on mRNAs with overly distant PA-signal can induce CSH(Stroup & Ji, 2023). Additionally, emerging evidences indicate a direct association of APA with the CSH and the gene expression(Stroup & Ji, 2023). Further analysis is required to shed light on the aspect of APA with CSH and its cellular implications.

Furthermore, the signals for the regulation of CSH during oxidative stress response appears to relay through Star-PAP in the CPA complex. Yet, the mechanism how PAP regulates cleavage reaction is still unclear. In vitro studies have shown requirement of PAP in the fractionated CPA components for the reconstitution of cleavage reaction (Christofori & Keller, 1989; Takagaki *et al*, 1988; Terns & Jacob, 1989). In the case of Star-PAP-mediated cleavage complex, Star-PAP directly binds the target mRNA and helps in the recruitment of CPSF components including CPSF73(Kandala *et al*., 2016; Laishram & Anderson, 2010). Nevertheless, cleavage efficiency was affected by the addition of PAP in the in vitro cleavage reaction (Christofori & Keller, 1989). Moreover, PAP activity inhibitor cordycepin addition inhibits the cleavage efficiency(Terns & Jacob, 1989). However, the exact mechanism how polyadenylation activity affects cleavage is still unclear. Our study suggests a potential relationship between the cleavage efficiency and the polyadenylation activity through CSH. Star-PAPs PRR region that splits the PAP domain regulates Star-PAP polyadenylation activity through phosphorylation particularly during the antioxidant response (Gonzales *et al*., 2008). Deletion of PRR on Star-PAP will compromise Star-PAPs anti-oxidant-mediated stimulation of cleavage complex assembly inducing CSH. This will inhibit the stimulation of cleavage and polyadenylation of target mRNAs. However, further studies are required to understand the actual role of the PRR region in regulating the fidelity of cleavage and how it transduces the stress signal via CSH of target mRNAs. It is likely mediated through phosphorylation by different kinases CKIα or PKCδ that phosphorylates Star-PAP(Gonzales *et al*., 2008; Li *et al*, 2012).

## MATERIALS AND METHODS

### In silico cleavage site analysis

We employed 3′-READS sequencing data (GSE84461) for the global CS usage for each PA-site in HEK293 cells(Li *et al*., 2017). For oxidative stress response, RNA-seq data for H_2_O_2_ treatment (GSE138290) in Jurkat cells and for sodium arsenide, (GSE101851) mouse data was employed for the analysis(Mullani N, 2021; Zheng *et al*., 2018). The raw data extracted (data had ∼7000000 reads per replicates) from the GEO database was quality checked and pre-processed using FastQC (http://www.bioinformatics.babraham.ac.uk/ projects/fastqc). RNA 3′-adapter sequence was identified from the FastQC and trimmed using cutadapt tool along with the PCR duplicate sequences. We obtained ∼3000000 trimmed reads that were further aligned to the human genome (hg38) using Bowtie2(Langmead & Salzberg, 2012). For the mouse data of sodium arsenate treatment, it was mapped to the mouse genome (mm9). The alignment output files in SAM/BAM format were used for subsequent analysis. The number of reads mapped were determined using samtools(Li *et al*, 2009). The uniquely mapped reads were then identified using the criteria for unique mapping with MAPQ score exceeding 10, along with the presence of at least two unaligned 5′ Ts indicative of the PA-tail was also considered. There were altogether ∼1725768 uniquely mapped reads with a mapping score of >10 having minimum of two 5′ Ts not aligned to the genome as described earlier(Li *et al*., 2017). The aligned data were then extracted based on CIGAR string using the python programming. Denotations of PA-sites were as described earlier(Li *et al*., 2017). It required a PA-site to be supported by a minimum of two PA-site supporting reads and contribute to over 5% of a gene′s expression as described earlier(Li *et al*., 2017). The filtered reads meeting the above criteria were then located within a 25-nucleotide range of cleavage site after the PA-signal were clustered for subsequent analyses. For cleavage site (CS), we considered the CS that has at least 5 supporting reads converging at the 3′-end of a transcript. Cleavage events occurring within 25 nucleotides downstream of a PA-signal hexamer were clustered as CS of a particular PA-site that has >5% of the total transcript abundance in at least one replicate samples. Formatting of data and processing were done using python/perl programming. Integrative genome viewer tool was used to visualise the chromosomal location of different CS of distinct mRNAs (Robinson *et al*, 2011). Circos plots were used to visualise the genome-wide distribution of CSH(Krzywinski *et al*, 2009). Expression of genes and PA-site was obtained from the total PA-sites reads for a gene or a specific PA-site and was determined by the counts of reads per million of a PA-site uniquely mapped in a sample as described earlier (Li *et al*., 2017). Relative expression of each PA-site was obtained dividing the counts per million reads of a PA-site with that of the total counts per million reads of the gene expressed relative to internal control *GAPDH*. Relative gene expression was plotted against the cleavage site heterogeneity using Matplotlib (Hunter, 2007).

### Cell lines, animals, transient transfections, and cellular treatments

Human Embryonic Kidney 293 (HEK293) and HeLa cells were obtained from the American Type Culture Collection. Cells were then cultured in Dulbecco’s modified Eagle’s medium (DMEM) (Himedia, IN) supplemented with high glucose in the presence of 10% foetal bovine serum and a mix of 50 IU/ml penicillin and streptomycin (Gibco, USA). Cell cultures were maintained in a CO_2_ incubator at 37°C (Thermo Scientific, USA).

For transfection, cells were cultured until they reached approximately 60% confluency, and transfections of siRNA oligos and plasmid DNA were carried out using a chemical method (calcium phosphate), as described earlier(Sudheesh *et al*., 2019). About 1.5 μg of plasmid DNA per 0.5 × 10^6^ cells or 80 nM of siRNA oligos (Eurogenetec, Belgium) were used for transfection. Transfection was repeated 24 hours later. Cells were collected 48 hours after transfection for downstream analysis of RNAs and proteins.

### Induction of anti-oxidant response

To induce oxidative stress response, HEK293 cells were grown to 70% confluency and treated with 100 μM tert-Butyl Hydroquinone (tBHQ) an oxidative stress agonist in DMSO (Sigma, St. Louis, Missouri, USA) for 4 hours, DMSO was added as the vehicle control(Sudheesh *et al*., 2019). To induce cells with peroxide stress 50 μM of H_2_O_2_ was added to the culture media, then incubated for 2 hours followed by harvesting as per the protocol described earlier(Mullani N, 2021). Heavy metal mediated oxidative stress was induced by treatment with 250 μM NaAsO_2_ in the cell culture media for 6 hours followed by harvesting the cells for further processing(Zheng *et al*., 2018).

### DNA Constructs

FLAG tagged pCMV Tag2a Reporter construct used in this study consists of *NQO1* coding sequence under the control of the pCMV promoter. Additionally, this construct is driven by the full-length *NQO1* 3′-UTR region approximately 1950 nucleotides long as described earlier(Sudheesh *et al*., 2019). The distal *NQO1* PA-site was used for the mutational analysis, this PA-site have five major cleavage sites that are consistently observed (namely P, S_1_, S_2_, S_3_, S_4_) downstream of which was mutated by site directed mutagenesis individually and in combination (see Supplemental Table S1). In the first batch of mutants one CS was mutated at a time namely *P^m^* (primary CS), *S^m^_1_* to *S^m^_4_* (subsidiary CS site S1to S4). Next batch of mutants has combinations of subsidiary CS mutations. To understand the characteristics of primary CS sequence and position, corresponding mutations of the CS (CA to GA, TA, AA) and position of the CS (-10 bp upstream, and downstream from the original CS position) were introduced(see Supplemental Table S1). Silent mutations to make the FLAG-NQO1 construct insensitive to the siRNA sequence employed for the NQO1 knockdown was introduced using pair of primers bearing three different mutations on the targeting sequence of the siRNA.

For the synthesis of in vitro transcribed *NQO1* 3′-UTR RNA, a UTR region (∼250 nucleotide) around the distal PA-site region of *NQO1* encompassing the sequences required for both cleavage and polyadenylation were cloned in pTZ vector (Sudheesh *et al*., 2019). Mutations of the CS were introduced in the pTZ19R construct, individual and in combination as described above. Primer sequences for the site directed mutagenesis are shown in Supplemental data.

### Site Directed Mutagenesis

Site directed mutagenesis of the CS or siRNA insensitive mutation were carried out as described earlier (Sudheesh *et al*., 2019)using a pair of primers incorporating the changed nucleotides. Briefly, around 50 ng of parent plasmid was taken and mutations were introduced by inverse PCR reaction using Pfu Ultra II polymerase (Agilent, cat no. 600674) enzyme using primer pairs having the respective mutations. The PCR reaction was digested using Dpn1(NEB, cat no. R0176L) to remove the template plasmid treatment for 1 hr at 37°C followed by transformation of DH5α competent cells with the in vitro amplified plasmid. Mutated plasmids were then confirmed by Sanger sequencing. Primers used for introducing mutations are listed in the Supplemental data.

### RNA immunoprecipitation

RNA IP was carried out as describe earlier (Mohanan *et al*, 2024). HEK293 cells transfected with wild-type FLAG-tagged NQO1 or various mutant variants were cross-linked with 1% formaldehyde for 10 minutes followed by quenching with 0.125 M glycine for 5 minutes. Cells were washed in 1 x PBS followed by lysis in 300 μl of cell lysis buffer (10 mM Tris-HCl pH 8.0, 10 mM NaCl, 0.2% NP-40, 1×EDTA-free Proteinase Inhibitor, and 1000 U RNase I, Promega, Madison, WI, USA). The crude nuclei were pelleted and resuspended in 400 μl nuclei lysis buffer (50 mM Tris-HCl pH 8.0, 10 mM EDTA, 1% SDS, 1× EDTA-free Proteinase Inhibitor, and 1000 U RNase I Promega, Madison, WI, USA). Supernatant obtained after sonication and centrifugation steps were digested with DNase I for 20 min at 37°C. The digestion was stopped by addition of 20 mM EDTA. Monoclonal antibodies against CPSF73, Star-PAP, and IgG control were added to the supernatant and incubated overnight at 4°C. Subsequently, Sepharose A beads equilibrated with IP dilution buffer were added, and the mixture was incubated for 2 hours at 4°C. The formed complexes were then pelleted, followed by washing in wash buffer for 3 times for 5 min each at 3500 rpm. It was then eluted with elution buffer (1% SDS and 100 mM Sodium bicarbonate, and proteinase K). Crosslinking was reversed by incubating the mixture at 67°C for 4 hours. RNA was isolated using Trizol, and cDNAs were synthesised using random hexamers and MMLV-RT (Invitrogen, Waltham, MA, USA). PCR amplification was carried out with gene-specific primers and products were visualised on an agarose gel.

### In vitro RNA synthesis

*NQO1* RNA substrate was derived from the plasmid pTZ-NQO1, which harbours the *NQO1* 3′-UTR under the T7 promoter. This plasmid includes a segment of the *NQO1* 3′-UTR containing the PA-signal, spanning from 120 base pairs upstream to 122 bp downstream of the cleavage site, inserted into the PstI and BamHI sites of the pTZ19R vector (Fermentas)(Sudheesh *et al*., 2019). Radiolabelled RNA substrates were uniformly prepared through in vitro transcription of linearised plasmid templates using the T7 transcription Kit (Invitrogen). A typical 20 μl reaction consisted of 1x reaction buffer (supplied in the kit), 20 units of RNase inhibitor, 1 μg of DNA, 500 μM each of ATP and CTP, 50 μM UTP, and 100 μM GTP, supplemented with 50 μCi of α-32P-UTP and 400 μM m7G cap analogue, along with 40 units of T7 RNA polymerase. Following transcription, RNA was purified using the Qiagen Mini RNeasy Kit for subsequent experimentation.

### In vitro pre-mRNA cleavage assay

Cleavage experiments were carried out as described earlier(Laishram & Anderson, 2010). Each reaction included a 32P-labeled pre-mRNA substrate in a 25 µl volume of cleavage buffer. The buffer contained 20 mM creatinine phosphate, 0.5 mM MgCl_2_, 10% glycerol, 10 mM HEPES (pH 7.9), 50 mM KCl, 0.05 mM EDTA, and 1% polyvinyl alcohol (PVA), supplemented with 0.8 mM 3′-dATP and 5 µl of HeLa nuclear extract. Reactions were incubated at 30°C for 2 hours and stopped by adding a proteinase K mixture comprising 2% sarcosyl, 100 mM Tris-Cl (pH 7.5), 20 mM EDTA, and 400 mg/ml proteinase K, followed by an additional incubation at 30°C for 10 minutes. RNA extraction was performed using phenol chloroform, followed by precipitation with absolute alcohol in the presence of 3 M ammonium acetate and 1 mg carrier tRNA. The extracted RNA was then analysed on a 6% urea denaturing gel.

### In vitro polyadenylation assay

In vitro polyadenylation assay was carried out as described earlier(Sudheesh *et al*., 2019). Briefly DNA templates with the forward T7 promoter and reverse primer ending with a CS was amplified from the pTZ19R *NQO1* 3′-UTR vector. The amplified templates were then subjected to in vitro RNA transcription using T7 RNA polymerase generating cleaved RNA templates, ending at different cleavage sites either the primary or the subsidiary sites of *NQO1*. RNA templates were incubated in an equivalent of ∼10 μg of HeLa nuclear extract prepared using the NEPER nuclear extraction kit (ThermoScientific) with α-32P ATP in a PAP assay buffer (250 mM NaCl, 50 mM Tris–HCl, and 10 mM MgCl_2_, pH 7.9) at 37°C for 1 hour. Then the polyadenylated RNA was precipitated and washed with 75% ethanol. RNA products were then analysed on a 6% urea-denaturing polyacrylamide gel and visualised by phosphor imaging.

### Immunoblotting experiment

For immunoblotting, cells were lysed in a cell lysis buffer comprising 0.06 M Tris-HCl (pH 7.4), 25% Glycerol, 2% SDS, 0.002% Bromophenol blue, and 1% β-mercaptoethanol (1× SDS-PAGE loading buffer) as described earlier (Shaji, 2024). The lysed samples were denatured by heating at 95°C for 20 minutes and then analysed on an SDS-PAGE gel in 1× Tris Glycine Buffer (25 mM Tris pH 8.0, 190 mM Glycine, 0.1% SDS). Following separation, the proteins were transferred onto a PVDF membrane using a transfer buffer (25 mM Tris-Cl pH 8.0, 190 mM Glycine, 20% Methanol) at 110 V for 90 minutes. The PVDF membrane was then blocked in 5% skimmed milk in 1× TBST (20 mM Tris pH 7.4, 150 mM NaCl, 0.1% Tween-20) for 45 minutes at room temperature (RT). Blots were subsequently incubated with primary antibodies at a dilution of 1:3000 in TBST. After three washes in TBST for 10 minutes each, blots were incubated with HRP-conjugated secondary antibody (Jackson Immuno Research Laboratory, West Grove, PA, USA) at a dilution of 1:5000. Finally, blots were imaged using a chemiluminescent substrate (Bio-Rad, Hercules, CA, USA) on an iBright FL1500 platform (Invitrogen, Waltham, MA, USA). A list of antibodies employed in the study is shown in Supplemental data.

### Immunoprecipitation experiment

For immunoprecipitation, cell lysates were sonicated (28% amplitude for 5 seconds on and off pulse for 5 minutes) in a cell lysis buffer as described earlier (Shaji, 2024) containing 100 mM KCl, 50 mM Tris-HCl (pH 7.4), 0.5% Nonidet NP-40, 5 mM EDTA, 200 mM NaVO_4_, 50 mM NaF, 50 mM L-glycerophosphate, and 1× EDTA-free protease inhibitor, supplemented with 100 μg/ml RNase A. Following sonication, the lysates were centrifuged at 12,500 rpm for 15 minutes at 4°C to collect the cellular lysates. For the FLAG tagged constructs of Star PAP lysates were incubated in the IP buffer equilibrated FLAG M2 beads overnight at 4°C. The following day, beads were centrifuged and precipitated, supernatant was discarded and washed three times with IP buffer for 10 minutes each. Finally, SDS loading buffer was added to the beads, and the mixture was loaded onto an SDS-PAGE gel after denaturation at 95°C for 5 minutes. The input loaded on the gel corresponded to 10% of the total protein utilized in each immunoprecipitation experiment(Shaji, 2024). Antibodies used for immunoblotting are listed in the Supplemental data.

### 3′-RACE and pre-mRNA cleavage assay

3′-RACE assays were conducted using 2 μg of total RNA extracted from HEK293 cells using an RNA isolation kit (Promega) following established protocols (Shaji, 2024). An engineered oligodT reverse primer (comprising 16 oligo dT nucleotides preceded by a 20-nucleotide engineered sequence at the 5′-end) was employed and reverse transcription reaction was performed with 100 units of MMLV-RT (Invitrogen, cat. 28,025–013) at 42°C for 1 hour, followed by inactivation at 70°C for 15 minutes. Subsequently, 250 ng of cDNA was amplified using a gene-specific forward primer (specific to each gene) and an AUAP reverse primer (specific to the engineered region of the oligo dT primer). The 3′-RACE products were then validated through sequencing. For in vivo cleavage assays, cleavage efficacy was evaluated using a pair of primers designed across the cleavage site, specifically amplifying the uncleaved pre-mRNA by qRT-PCR. The levels of uncleaved pre-mRNAs were normalised to internal control GAPDH levels and expressed relative to total mRNA levels (fold-change over total mRNA). Primers used for 3′-RACE and pre-mRNA cleavage assay are listed in the Supplemental data.

### Real-Time qPCR

The quantitative real-time PCR (qRT-PCR) experiments were conducted using the CFX98 multi-color system (Bio-Rad). iTaq universal SYBR Green Supermix was utilised for the PCR reactions with cDNA synthesised by reverse transcription using MMLV-RT from 2 μg total RNA samples as described earlier(Shaji, 2024). Melt curve analysis was performed for each qRT-PCR experiment to verify single product amplification. To ensure specificity and accuracy of the PCR reaction, primer efficiency was maintained close to 100% in all experiments.

For quantification of the real-time experiments, target mRNA levels were normalised to GAPDH levels, and changes in expression levels were expressed as fold change relative to the control samples using the Pfaffl method. Three independent experiments were conducted for each qPCR experiment (n > 3). Primers employed in the qRT-PCR are shown in Supplemental data.

### MTT assay

The MTT assay, was carried out to assess cell viability as described earlier(Mohanan *et al*., 2024). Transfected cells were seeded onto a 96-well plate at a density of 1 × 10^4^ cells per well. Following a 48-hour incubation period in a humidified incubator at 37°C with 5% CO2, 3-(4,5-dimethylthiazolyl-2-yl)-2–5 diphenyl tetrazolium bromide (MTT, Sigma, St. Louis, Missouri, USA) was introduced. Subsequently, the plates were further incubated for 3 hours, followed by the addition of isopropanol subsequent to removal of the culture medium. The resultant formazan crystals were resuspended and absorbance was measured at 570 nm utilizing a Varioskan LUX multimode microplate reader (ThermoScientific).

### Trypan Blue Staining

Trypan blue solution was prepared by mixing equal parts of 0.4% trypan blue solution and cell suspension followed by incubation for 3 minutes at room temperature. Then, a drop of the trypan blue/cell mixture was applied onto a haemocytometer and the unstained (viable) and stained (nonviable) cells were counted separately using haemocytometer. To determine the total number of cells of the aliquot, counts of viable and nonviable cells were added together and the dilution factor multiplied. Finally, the percentage of viable cells were calculated in each treatment condition(Strober, 2015).

### Catalase activity assays

Catalase activity determination was performed using Catalase Assay Kit (Cayman chemical cat no. 707002). Briefly, cells were collected by cell scrapper and then homogenised in 1 ml cold lysis buffer (50 mM potassium phosphate, pH 7, containing 1 mM EDTA). Then, the cell suspension was centrifuged at 10,000 g for 15 min at 4° C and then the supernatant was collected. Further, a series of catalase standards was prepared with known concentrations (from 0 to 10 nmol/ml) using a catalase standard solution. Samples were transferred to a microplate well and then 20 μl hydrogen peroxide solution was added to all wells including standards to initiate the reaction. The microplate was incubated at room temperature for 20 minutes to allow the catalase enzyme to react with hydrogen peroxide. Then, 30 μl of potassium hydroxide was added to each well to terminate the reaction followed by 30 μl of chromogen. The plate was further incubated for 10 min at room temperature in a shaker and then 10 µl of catalase potassium periodide added to each well followed by an additional incubation in the dark for 5 min and absorbance was measured at 540 nm using spectrophotometer.

### RNA biotinylation and pulldown

Biotinylated RNA was prepared by in vitro transcription on the pTZ19R plasmid template with Biotin UTP instead of radiolabelled UTP in the in vitro RNA synthesis method. For biotinylation pull down, HEK cell lysates were prepared from 90% confluent 10 cm dish in lysis buffer (10 mM HEPES, pH-7, 200 mM NaCl, 1% Triton-X 100, 10 mM MgCl_2_, 1 mM DTT and protease inhibitor) as described earlier(Francis & Laishram, 2021). Briefly, 30 microgram of biotinylated RNA was incubated with 60 ml of pre-washed avidin-agarose beads at 4°C for 1 hour. Agarose beads were washed three times with RNA binding buffer (10 mM Tris HCl, 0.1 M KCl and 10 mM MgCl_2_), incubated with 5 mg total protein equivalent cell lysates overnight at 4°C. Agarose beads were collected, washed and proteins were eluted using SDS loading buffer after boiling for 10 minutes at 95°C. Associated proteins were analysed by Western blotting using specific antibodies.

### Blot quantification

Quantification of blots was performed using Image J software (NIH)(Schneider *et al*, 2012). Three replicates of blots were analysed to quantify relative intensities. Bands were quantified in arbitrary units and expressed relative to a reference protein. In the case of immunoprecipitation, bands were normalised to IgG and expressed relative to the Input sample.

### Statistics

Statistical significance was determined using ANOVA to analyse the differences in the means of experimental replicates from each experiment using GraphPad Prism 10. A minimum of three independent experiments were utilised for quantification of each experimental dataset. All data are presented as mean ± standard error mean (SEM). Differences were considered statistically significant at a p-value < 0.05.

## Acknowledgement

We thank Fiona P. Ukken (University of Maryland, USA) and RSL lab members for carefully reading the manuscript.

## Data Availability Statement

The article and supplemental data contain all the data generated during this study.

## Author Contribution

FS carried out all experiments in the paper. JA contributed to bioinformatic analysis. RSL planned and conceptualised the work and directed the overall project.

